# From linear to nonlinear gait measures: validity of a multi-view markerless motion capture system

**DOI:** 10.64898/2026.09.07.749073

**Authors:** Baptiste Perthuy, Hugues Vinzant, Clément Brifault, Nicolas Lefèvre, Alexandre Dalibot, Sofiane Ramdani, Leslie M. Decker

## Abstract

Markerless motion capture offers a practical alternative to marker-based optoelectronic systems, yet validation studies have focused almost exclusively on linear gait measures. Nonlinear measures of gait dynamics, sensitive to the fine temporal structure of locomotor signals, remain unvalidated in markerless systems. This study assessed the concurrent validity of a three-camera markerless system against a 15-camera optoelectronic system across four treadmill speeds in 22 healthy adults. Inter-system agreement was evaluated for spatiotemporal parameters (mean, variability, detrended fluctuation analysis [DFA] scaling exponents) and for joint angle and trunk acceleration time series (maximum Lyapunov exponents, sample entropy, Attractor Complexity Index [ACI]). Temporal measures reached near-perfect agreement, and spatial measures showed good to excellent agreement with small speed-dependent positive biases. Sagittal-plane joint kinematic waveforms were compared using statistical parametric mapping, with root-mean-square error (RMSE) reported for significant intervals. Agreement was best at the hip, with knee and ankle showing comparable, higher error (RMSE: 1.2–3.0° hip, 3.7–5.5° knee and 3.8–5.1° ankle). ACI demonstrated moderate to good agreement across all joints and good agreement across trunk acceleration directions. Most DFA scaling exponents for step-based series supported group-level comparisons: both measures are usable for markerless assessment of gait’s nonlinear dynamics. Maximum Lyapunov exponents and sample entropy showed lower absolute agreement: at the hip and knee, they preserved inter-individual ranking and remained usable for within-system group comparisons, but agreement collapsed at the ankle and for trunk sample entropy, indicating these measures still need refinement. These findings define a tiered, measure-specific scope for markerless gait analysis, extending validation beyond spatiotemporal parameters.

**Graphical abstract:** Concurrent validity of a three-camera markerless motion capture system against a 15-camera optoelectronic reference during treadmill walking

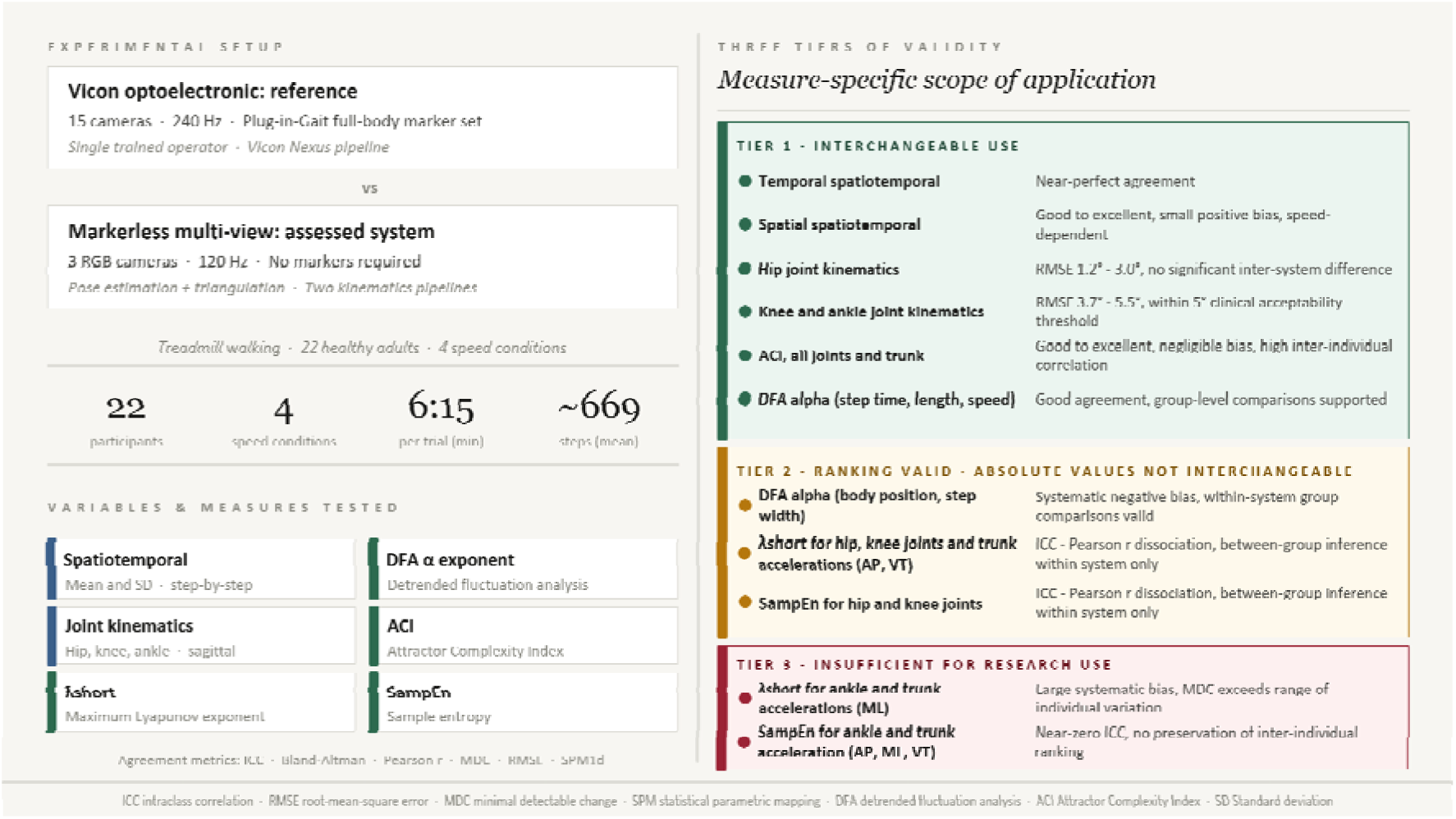

## 1. INTRODUCTION

Quantitative analysis provides a sensitive framework for identifying locomotor impairment, characterizing disease-related motor changes, and monitoring intervention effects across neurological, musculoskeletal, and aging-related conditions (Baker, 2006; Muro-de-la-Herran et al., 2014). Marker-based optoelectronic systems remain the laboratory reference standard for three-dimensional spatiotemporal and kinematic measurements but require costly, dedicated infrastructure and trained operators for consistent marker placement, and remain susceptible to soft-tissue artifact (Chiari et al., 2005; Gorton et al., 2009; Peters et al., 2010).

Markerless motion-capture systems based on video pose estimation and three-dimensional reconstruction offer a practical alternative to marker-based systems. They extend quantitative gait analysis beyond specialized laboratories by removing physical markers. Recent validation studies report good agreement with marker-based systems for spatiotemporal parameters and sagittal-plane lower-limb kinematics, although agreement remains lower at the ankle than at the hip or knee (Kanko, Laende, Davis, et al., 2021a; Scataglini et al., 2024).

Markerless validation studies have nonetheless focused almost exclusively on linear gait measures, namely summary statistics of mean locomotor output and step-to-step variability (Hausdorff, 2005; Hollman et al., 2011). Such measures capture the gait pattern and its step-to-step fluctuation amplitude but reveal little about how gait is regulated from one step to the next (Stergiou & Decker, 2011). Nonlinear analyses, including the divergence of movement trajectories, sample entropy, and long-range temporal correlations, quantify the temporal structure of these fluctuations, reflecting stability, complexity, and adaptive control (Bruijn et al., 2013; Cignetti et al., 2011; Hausdorff, 2005; Yentes & Raffalt, 2021).

This distinction matters for validation. Unlike linear measures, nonlinear measures depend critically on the fine temporal structure of the signal, making their estimation sensitive to measurement noise, filtering strategy, and other preprocessing choices (Mehdizadeh & Sanjari, 2017; Raffalt et al., 2020; Yentes & Raffalt, 2021). Good agreement for the linear measures therefore does not guarantee comparable validity for the nonlinear ones. Because nonlinear analyses require long, stationary recordings, treadmill walking provides an ideal setting to assess them, and this assessment has not yet been performed for recent video-based markerless systems.

The present study aimed to validate a multi-view markerless motion-capture system against a marker-based optoelectronic reference during treadmill walking. Inter-system agreement was evaluated for (i) linear measures of conventional gait variables, namely spatiotemporal parameters and sagittal-plane joint kinematics, and (ii) nonlinear measures indexing the temporal structure of gait fluctuations. Agreement was hypothesized to be strong for the linear measures but not necessarily to extend to the nonlinear ones, given their sensitivity to the fine temporal structure of the signal.

## 2. METHODS

### 2.1. Participants

Twenty-two healthy adults (9 women, 13 men; age 29.4 ± 10.4 years, height 1.75 ± 0.10 m, mass 68.7 ± 12.3 kg) with no neurological, musculoskeletal, or vestibular disorders and no lower-limb injury in the preceding six months were enrolled. All provided written informed consent (ethics approval: CERSTAPS, IRB00012476-2024-06-09-338).

### 2.2. Experimental protocol

Participants completed four 6 min 30 s walking trials on an instrumented dual-belt treadmill (M-Gait, Motekforce Link, Netherlands). Preferred walking speed (PWS) was determined using a standardized incremental protocol (Jordan et al., 2007). A self-paced trial was always performed first, followed by three fixed-speed trials at 100%, 120%, and 140% PWS in randomized order.

### 2.3. Instrumentation and data acquisition

Three-dimensional motion data were acquired simultaneously from a 15-camera optoelectronic system (Vicon Motion Systems, Oxford, UK, 240 Hz) with retroreflective markers placed by a single trained operator according to the Plug-in-Gait Full Body marker set (Davis et al., 1991), and a markerless system comprising three RGB cameras (iPhone 17 Pro, Apple Inc., Cupertino, CA, USA, 120 Hz). For the markerless system, a three-camera multi-view configuration was chosen over a single camera because triangulating keypoints across synchronized views reduces self-occlusion and three-dimensional reconstruction error relative to monocular estimation (Horsak et al., 2025). The three cameras were positioned to keep the sagittal-plane landmarks continuously visible throughout the gait cycle: one behind the treadmill along the sagittal plane and two lateral cameras at approximately 45° from that plane (D’Souza et al., 2024; Stenum et al., 2021). Camera intrinsics were estimated once using dynamic 11 x 9 black-and-white checkerboard calibration, and extrinsics were re-estimated before each session with the same checkerboard. For each camera view, a bounding box person detector (RTMDet) and a two-dimensional pose estimator (RTMPose, large model) – selected for its state-of-the-art accuracy among open-source real-time 2D pose estimators (Jiang et al., 2023) – generated frame-wise keypoints based on a Halpe-26 skeleton definition. Three-dimensional keypoints were then reconstructed by triangulation. Systems were synchronized in two steps: a hardware trigger provided coarse, frame-level alignment, and the residual sub-frame temporal offset between systems was then estimated and corrected by maximizing the cross-correlation of a common signal. All trajectories were low-pass filtered with a fourth-order zero-lag Butterworth filter at 8 Hz (Crenna et al., 2021), Vicon data were downsampled from 240 Hz to 120 Hz, and each trial was cropped to a common 6 min 15 s steady-state segment.

### 2.4. Gait event detection and segmentation

Heel-strike and toe-off events were detected identically in both systems using prominence-based peak detection on anteroposterior heel and toe trajectories (Zeni et al., 2008). The number of analyzed events was matched within each participant-by-condition pair to ensure valid inter-system comparison, yielding a mean of 669 ± 69 analyzed steps.

### 2.5. Spatiotemporal parameters

Spatiotemporal parameters were computed on a step-by-step basis, except for phase-related variables such as single-support and double-support times, which were expressed as percentages of the stride cycle. **Supplementary Table S1** summarizes all extracted spatiotemporal parameters with their definitions and formulas.

### 2.6. Joint kinematics

Sagittal-plane flexion-extension angles of the hip, knee, and ankle were extracted and time-normalized to 101 points per gait cycle. Vicon angles were obtained from the Plug-in-Gait pipeline implemented in Vicon Nexus (version 2.16, Vicon Motion Systems, Oxford, UK). Markerless angles were derived using two approaches. The first used Pose2Sim (Pagnon et al., 2021, 2022b, 2022a), an open-source markerless pipeline solving inverse kinematics via the OpenSim musculoskeletal modeling framework (Delp et al., 2007) with the LaiUhlrich2022 model. The second computed ZYX Euler angles directly from segment vectors defined by anatomical keypoints, retaining only the flexion-extension component as a lighter alternative not relying on musculoskeletal model assumptions.

### 2.7. Acceleration signals

A virtual trunk point was defined as the midpoint between C7 and the pelvic center in the Vicon system, and as the midpoint between the neck and mid-hip keypoints in the markerless system. Markerless position trajectories were first de-spiked with a Hampel filter (window = 7 samples, threshold = 3) to remove the isolated outliers characteristic of pose-estimation reconstruction (Pagnon et al., 2022b), and then low-pass filtered with a zero-lag fourth-order Butterworth filter (6 Hz cut-off), a standard choice for gait kinematics prior to differentiation (Winter, 2009). Vicon trajectories were filtered with the same zero-lag Butterworth filter but without the preceding Hampel step, as they were free of spike-like reconstruction artifacts. Accelerations were then obtained in both systems by fourth-order Savitzky-Golay differentiation with a 21-sample window (Savitzky & Golay, 1964). **Figure 1** shows representative trunk position, velocity, and acceleration signals from both systems, illustrating the effect of successive differentiation on the markerless signal.

**Figure 1.**
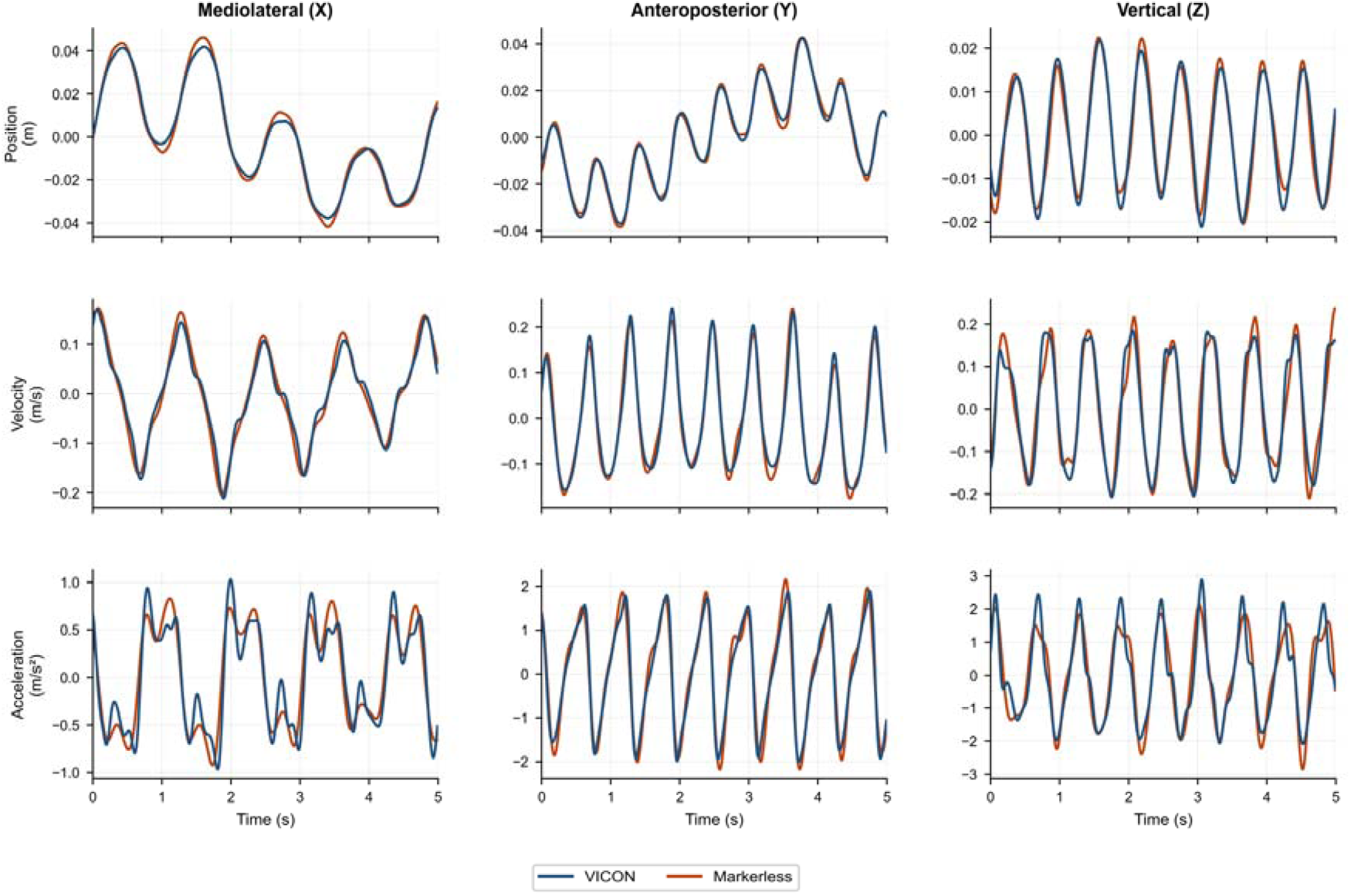
Five seconds segments of trunk position, velocity, and acceleration signals from both systems.

**Figure 2.**
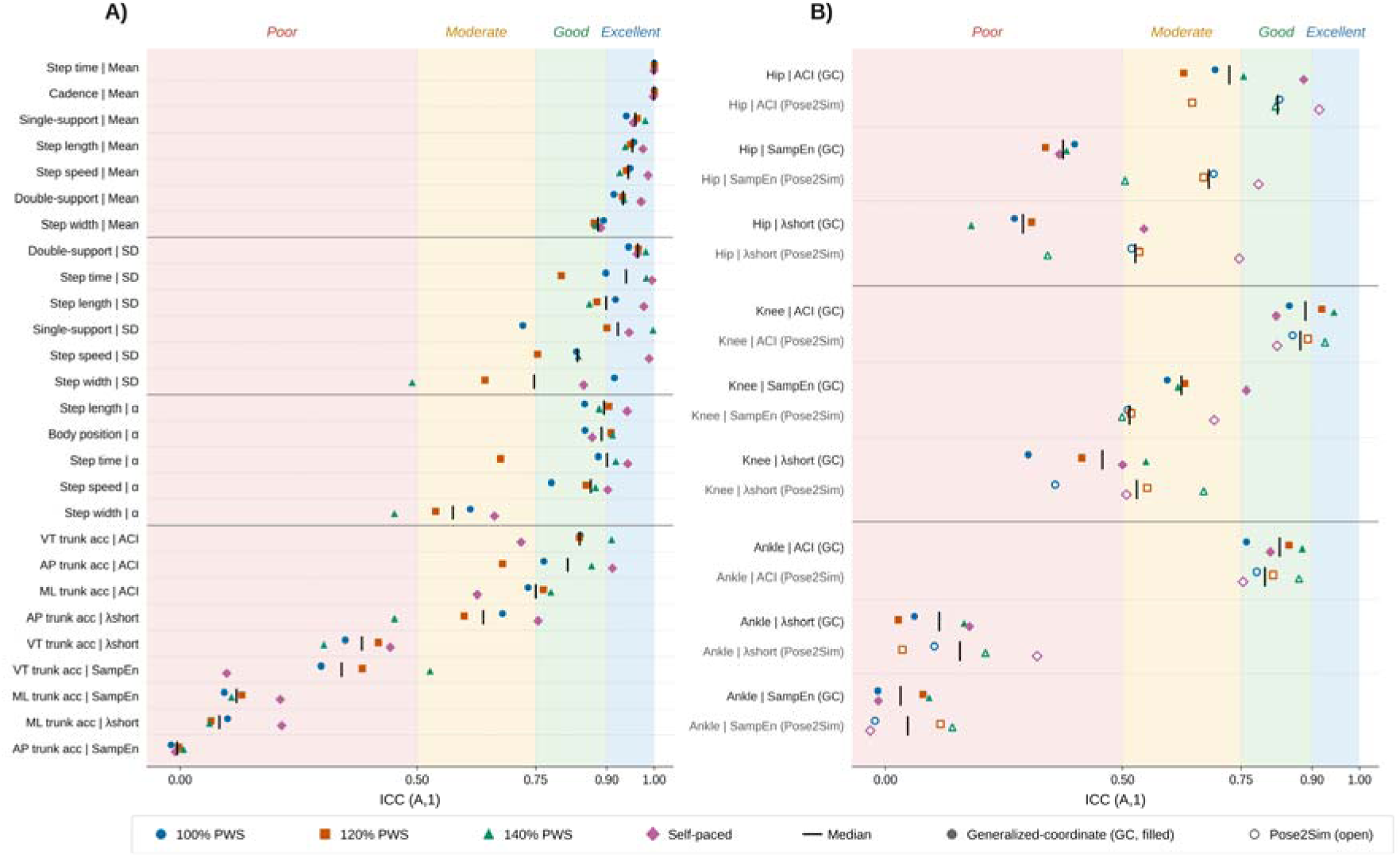
Inter-system agreement (ICC(A,1)) across all gait measures and walking speed conditions. **A)** Spatiotemporal parameters (means, standard deviations, DFA scaling exponents) and nonlinear trunk acceleration metrics. **B)** Nonlinear dynamics of sagittal-plane joint angle trajectories for the generalized-coordinate (GC, filled markers) and Pose2Sim (open markers) approaches. Each marker represents the ICC for a single walking speed condition, and the vertical bar indicates the median across conditions. Background shading denotes ICC interpretation bands: poor (< 0.50), moderate (0.50-0.75), good (0.75-0.90), and excellent (> 0.90) (Koo & Li, 2016). Measures are ordered by median ICC within each domain. SD: standard deviation. λshort: maximum Lyapunov exponent. ACI: attractor complexity index. SampEn: sample entropy.

### 2.8. Outcome measures: linear and nonlinear analysis of step-based and continuous gait signals

Linear measures consisted of the mean and standard deviation (SD) of each step-by-step time series per condition. For discrete step-based series, long-range temporal correlations were assessed using detrended fluctuation analysis (DFA), which yields a scaling exponent α (Peng et al., 1995). Values above 0.5 indicate persistent correlations, where deviations tend to propagate in the same direction across successive steps, while values below 0.5 indicate antipersistence, reflecting tighter step-to-step error correction consistent with stronger regulation of timing and foot placement under treadmill speed and position constraints. For continuous signals including sagittal-plane joint angles and trunk accelerations in the anteroposterior, mediolateral, and vertical directions, three complementary metrics were computed, each capturing a distinct aspect of the temporal structure of the time series (Stergiou, 2016; Stergiou & Decker, 2011). Divergence of movement trajectories was quantified using the short-term maximum Lyapunov exponent (λshort, Rosenstein et al., 1993), where larger values indicate greater sensitivity to small perturbations and reduced local stability. Signal regularity was assessed using sample entropy (SampEn, Richman & Moorman, 2000), where lower values reflect more regular and predictable fluctuations. Long-range dynamical complexity was characterized using the Attractor Complexity Index (ACI, Terrier, 2019), where higher values reflect richer temporal organization and lower values indicate a simplification of locomotor dynamics. To quantify how differentiation affects these estimates, the three continuous-signal metrics were additionally applied to the trunk position and velocity signals from which the accelerations were derived (**Supplementary Tables S7 and S8**). All preprocessing steps and state-space reconstruction parameters are detailed in **Supplementary Appendix A**.

### 2.9. Statistical analysis

Inter-system agreement was assessed per condition using: two-way mixed-effects, single-measure intraclass correlation coefficient for absolute agreement (ICC(A,1), interpreted per Koo & Li, 2016), Pearson’s r, Bland-Altman bias with 95% limits of agreement (Bland & Altman, 1986), proportional bias regression, standard error of measurement (SEM), and the minimal detectable change (MDC). Agreement statistics are reported in the main text as median [min, max] across conditions, and MDC as its maximum value (MDCmax). Full condition-specific results are in the **Supplementary Materials**. Speed-condition effects on inter-system differences (markerless minus Vicon) were assessed by one-way repeated-measures ANOVA (four levels), with Greenhouse-Geisser correction where needed and Benjamini-Hochberg FDR correction across the 45 measures (pFDR < 0.05). Significant effects were followed by Bonferroni-adjusted pairwise comparisons. Partial eta squared (η²p) was reported as the effect size. For joint kinematics, waveform-level differences were tested with paired SPM{t} (SPM1d, Pataky, 2012, α = 0.05), and the local RMSE was reported when significant intervals were found. The overall RMSE was also computed by combining the data from all strides and participants.

## 3. RESULTS

### 3.1. Agreement of linear analysis of step-based and continuous gait signals

#### 3.1.1. Mean and fluctuation amplitude of spatiotemporal parameters

Temporal variables showed near-perfect concordance across all walking speeds (**Table 1**, **Supplementary Table S2**). Step time and cadence yielded ICC = 1.000 at every condition, and double-support and single-support times reached median ICCs of 0.934 and 0.960, respectively, with negligible biases (≤ 0.001 s). Spatial variables showed good to excellent agreement, with small positive biases for step speed (median ICC = 0.945, bias = 0.027 m/s) and step length (ICC = 0.954, bias = 0.014 m), indicating slight markerless overestimation. Step width showed the lowest agreement (ICC = 0.881). Pearson’s r consistently exceeded ICC (e.g., r = 0.988 for step speed), confirming that inter-individual ranking was preserved despite the systematic offset.

**Table 1.** Inter-system agreement for mean spatiotemporal parameters, summarized across walking speed conditions.

| Measure | ICC(A,1)<br>median [min , max] | r<br>median [min, max] | Bias<br>median [min, max] | MDCmax |
| --- | --- | --- | --- | --- |
| Step time (s) | 1.000 [1.000, 1.000] | 1.000 [1.000, 1.000] | 0.000 [0.000, 0.000] | 0.000 |
| Cadence<br>(steps/min) | 1.000 [1.000, 1.000] | 1.000 [1.000, 1.000] | -0.003 [-0.004, 0.000] | 0.011 |
| Single-support<br>time (s) | 0.960 [0.941, 0.981] | 0.960 [0.940, 0.982] | -0.001 [-0.001, 0.000] | 0.015 |
| Step speed (m/s) | 0.945 [0.927, 0.987] | 0.988 [0.984, 0.998] | 0.027 [0.021, 0.037] | 0.078 |
| Step length (m) | 0.954 [0.939, 0.976] | 0.990 [0.986, 0.996] | 0.014 [0.012, 0.018] | 0.038 |
| Double-support<br>time (s) | 0.934 [0.915, 0.972] | 0.947 [0.914, 0.972] | 0.001 [-0.000, 0.002] | 0.031 |
| Step width (m) | 0.881 [0.873, 0.893] | 0.913 [0.902, 0.927] | 0.007 [0.006, 0.007] | 0.023 |

**Notes:** Each cell summarizes four walking speed conditions (100%, 120%, 140% of preferred walking speed, and self-paced). ICC(A,1): two-way mixed-effects, absolute agreement, single measure. Bias: mean inter-system difference (markerless, Vicon). r: Pearson's correlation. MDCmax: minimal detectable change at 95% confidence, maximum across conditions. Full condition-specific results, 95% limits of agreement, proportional bias tests, and Bland-Altman plots are provided in **Supplementary Materials**.

Step-to-step variability followed a similar hierarchy (**Table 2**, **Supplementary Table S3**), with most measures reaching good to excellent agreement (median ICC from 0.838 to 0.965). Step width SD was the exception (ICC = 0.489 - 0.916 across speeds), with significant proportional bias at faster speeds.

**Table 2.** Inter-system agreement for step-to-step variability (standard deviation) of spatiotemporal parameters, summarized across walking speed conditions.

| Measure | ICC(A,1)<br>median [min , max] | r<br>median [min, max] | Bias<br>median [min, max] | MDCmax |
| --- | --- | --- | --- | --- |
| Step time SD (s) | 0.941 [0.804, 0.995] | 0.942 [0.869, 0.995] | 0.000 [0.000, 0.001] | 0.003 |
| Step speed SD<br>(m/s) | 0.838 [0.756, 0.989] | 0.907 [0.865, 0.994] | 0.004 [0.003, 0.004] | 0.013 |
| Step length SD<br>(m) | 0.899 [0.863, 0.978] | 0.944 [0.911, 0.989] | 0.002 [0.002, 0.002] | 0.005 |
| Step width SD<br>(m) | 0.747 [0.489, 0.916] | 0.804 [0.560, 0.928] | 0.002 [0.001, 0.002] | 0.008 |
| Double-support<br>time SD (s) | 0.965 [0.946, 0.982] | 0.966 [0.945, 0.983] | -0.001 [-0.001, 0.000] | 0.010 |
| Single-support<br>time SD (s) | 0.923 [0.723, 0.997] | 0.929 [0.786, 0.997] | 0.000 [0.000, 0.001] | 0.003 |

**Notes:** See **Table 1**. SD: standard deviation.

#### 3.1.2. Sagittal-plane joint kinematic waveforms

Agreement was best at the hip, with knee and ankle showing comparable, higher error. The hip showed the lowest overall RMSE (generalized-coordinate: 1.2° to 1.8°, Pose2Sim: 2.1° to 3.0°), with no significant inter-system difference at any speed. The knee showed intermediate overall RMSE (3.7° to 4.2° and 4.4° to 5.5°, respectively), with both approaches producing a positive mean bias (3.1° to 4.9°) and significant differences concentrated during loading response and swing. The ankle showed overall RMSE of 4.0° to 4.3° (generalized-coordinate) and 3.8° to 5.1° (Pose2Sim), with the two approaches displaying opposite bias directions. Significant inter-system differences and local RMSE values are detailed in **Figure 3**.

**Figure 3.**
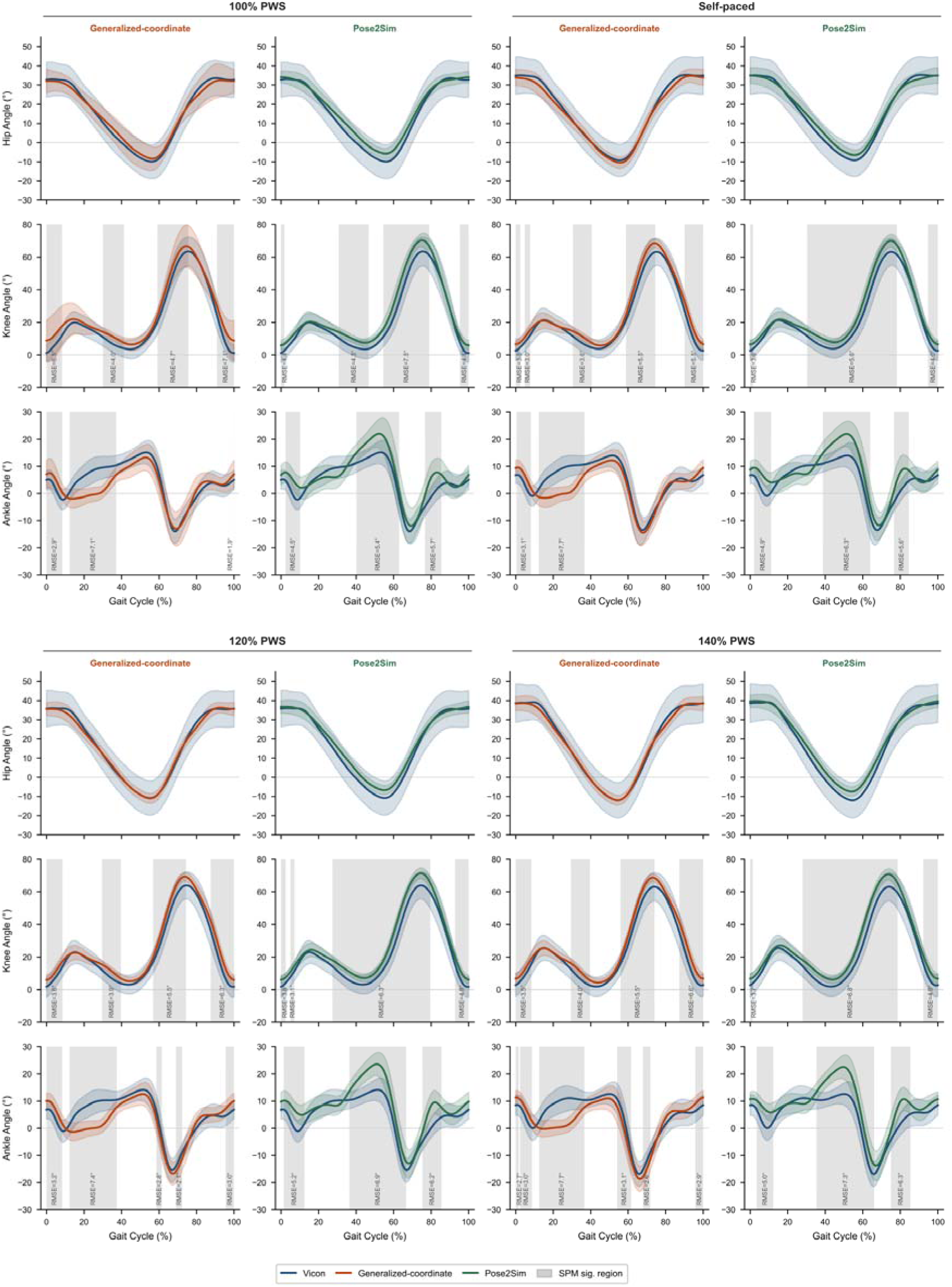
Sagittal-plane joint angle waveforms across walking speed conditions. Each panel shows group mean trajectories (solid lines) ± 1 SD (shaded areas) for Vicon (blue), the generalized-coordinate method (orange), and Pose2Sim (green). Gray regions indicate portions of the gait cycle with statistically significant inter-system differences (paired SPM{t}, α = 0.05), with local RMSE values reported for each significant cluster.

### 3.2. Agreement of nonlinear analysis of step-based and continuous gait signals

#### 3.2.1. Temporal correlations of step-based spatiotemporal parameters

DFA scaling exponents showed generally lower agreement than the corresponding linear statistics (**Table 3**, **Supplementary Table S4**). Step time α, step length α, and step speed α showed good agreement (median ICC = 0.901, 0.894, and 0.866, respectively), with small negative biases indicating marginally lower scaling exponents from the markerless system. Body position α showed a dissociation between ICC (median = 0.889) and Pearson’s r (median = 0.973) with a systematic negative bias (median = −0.068), indicating consistent underestimation of mediolateral temporal persistence while preserving inter-individual ranking. Step width α showed the lowest agreement (median ICC = 0.576, range 0.452 to 0.663), with all conditions falling in the moderate-to-poor range and a systematic negative bias (median = −0.062).

**Table 3.** Inter-system agreement for temporal correlations (DFA scaling exponent α) of step-based parameters, summarized across walking speed conditions.

| Measure | ICC(A,1)<br>median [min, max] | r<br>median [min, max] | Bias<br>median [min, max] | MDCmax |
| --- | --- | --- | --- | --- |
| Step time $\alpha$ | 0.901 [0.676, 0.944] | 0.905 [0.772, 0.943] | -0.012 [-0.026, -0.008] | 0.116 |
| Step length $\alpha$ | 0.894 [0.853, 0.943] | 0.894 [0.854, 0.953] | -0.014 [-0.025, -0.008] | 0.137 |
| Step speed $\alpha$ | 0.866 [0.793, 0.902] | 0.873 [0.794, 0.902] | -0.012 [-0.022, -0.006] | 0.169 |
| Body position $\alpha$ | 0.889 [0.854, 0.912] | 0.973 [0.952, 0.983] | -0.068 [-0.087, -0.057] | 0.185 |
| Step width $\alpha$ | 0.576 [0.452, 0.663] | 0.701 [0.552, 0.739] | -0.062 [-0.074, -0.061] | 0.241 |

**Notes:** See **Table 1**

#### 3.2.2. Nonlinear dynamics of joint angle trajectories

Agreement for joint-level nonlinear measures is summarized in **Table 4**, with condition-specific results for both the generalized-coordinate and Pose2Sim methods reported in **Supplementary Table S5**.

**Table 4.**
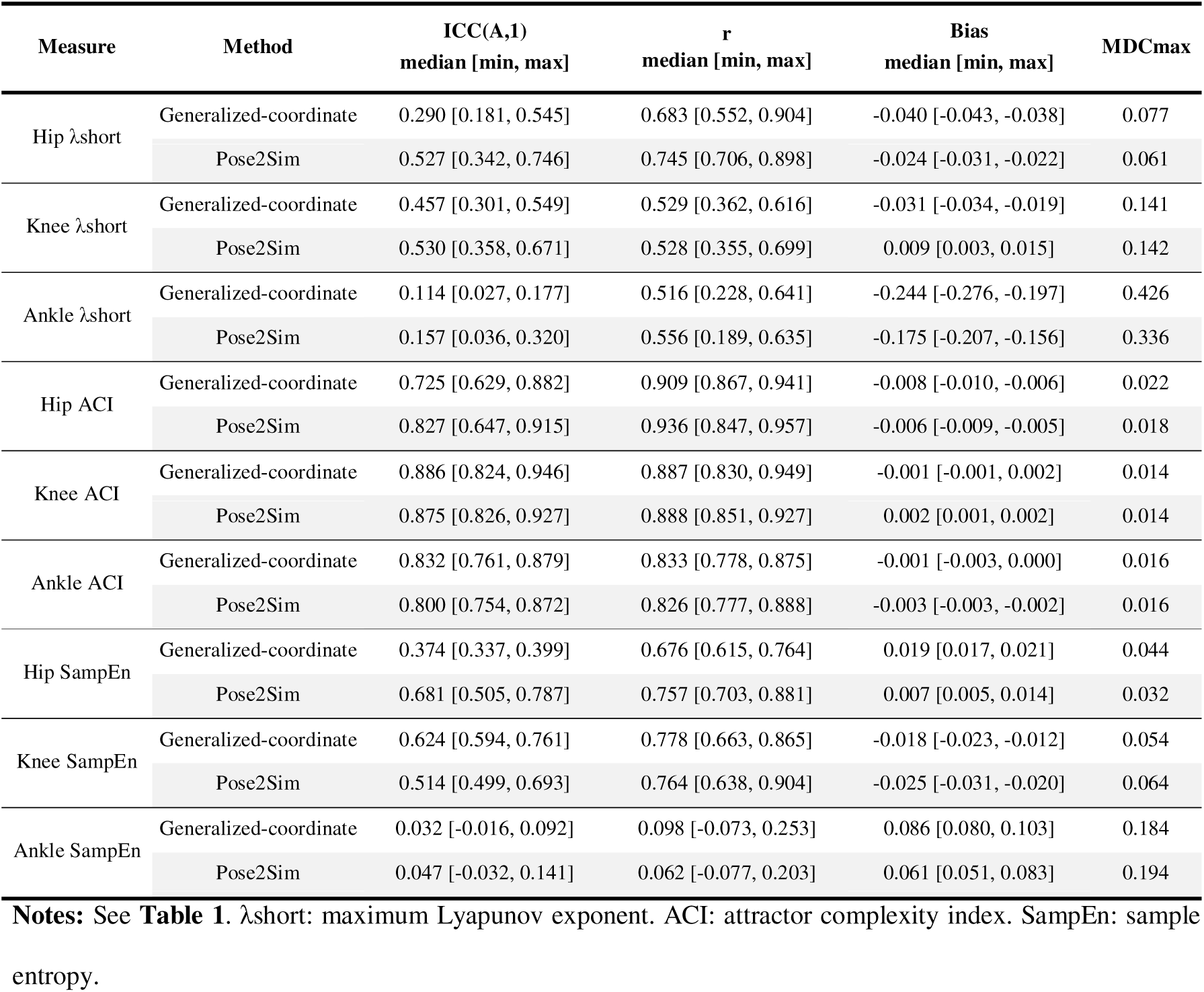
Inter-system agreement for nonlinear measures of sagittal-plane joint angle trajectories, summarized across walking speed conditions. Results are shown for the generalized-coordinate and Pose2Sim methods.

ACI showed the strongest agreement among the three joint-level nonlinear metrics. Knee ACI reached good to excellent agreement with both methods (generalized-coordinate: ICC = 0.886, Pose2Sim: ICC = 0.875), with negligible biases and high correlations (r ≥ 0.887). Ankle ACI showed similar performance (ICC = 0.832 and 0.800). Hip ACI was slightly lower (ICC = 0.725 and 0.827) with a small negative bias (−0.006 to −0.008) and high correlations (r ≥ 0.847), indicating a systematic offset without loss of inter-individual ranking.

λshort showed substantially lower agreement with a proximal-to-distal gradient. Hip λshort reached poor to moderate agreement (generalized-coordinate: ICC = 0.290, Pose2Sim: ICC = 0.527) with systematic negative biases. Knee λshort was moderate with both approaches (ICC = 0.457 and 0.530). Ankle λshort was poor across all conditions (ICC = 0.114 and 0.157), with large negative biases (median −0.244 and −0.175) and MDCmax > 0.3, indicating inter-system discrepancy exceeded the range of meaningful individual differences. For all joints, Pearson’s r exceeded ICC, indicating better preservation of inter-individual ranking than absolute values.

SampEn agreement varied across joints and methods. Hip SampEn was poor with the generalized-coordinate method (ICC = 0.374) but moderate with Pose2Sim (ICC = 0.681), with a positive bias in both cases. Knee SampEn showed moderate agreement with both methods (ICC = 0.624 and 0.514), with a negative bias. Ankle SampEn showed no meaningful agreement with either method (ICC = 0.032 and 0.047), with near-zero correlations and large positive biases, indicating failure to reproduce between-participant differences.

#### 3.2.3. Nonlinear dynamics of trunk accelerations

Agreement for trunk-level nonlinear measures is summarized in **Table 5**, with condition-specific results in **Supplementary Table S6**.

**Table 5.** Inter-system agreement for nonlinear measures of trunk accelerations, summarized across walking speed conditions.

| Measure | ICC(A,1)<br>median [min, max] | r<br>median [min, max] | Bias<br>median [min, max] | MDCmax |
| --- | --- | --- | --- | --- |
| AP trunk acc. $\lambda_{\text{short}}$ | 0.639 [0.452, 0.755] | 0.641 [0.602, 0.750] | 0.019 [0.010, 0.084] | 0.222 |
| ML trunk acc. $\lambda_{\text{short}}$ | 0.083 [0.062, 0.214] | 0.581 [0.533, 0.720] | 0.371 [0.334, 0.467] | 0.701 |
| VT trunk acc. $\lambda_{\text{short}}$ | 0.383 [0.303, 0.443] | 0.797 [0.694, 0.818] | 0.184 [0.166, 0.189] | 0.365 |
| AP trunk acc. ACI | 0.818 [0.680, 0.912] | 0.899 [0.820, 0.929] | -0.005 [-0.007, -0.003] | 0.016 |
| ML trunk acc. ACI | 0.750 [0.627, 0.782] | 0.757 [0.717, 0.775] | -0.001 [-0.002, 0.004] | 0.018 |
| VT trunk acc. ACI | 0.843 [0.719, 0.910] | 0.925 [0.857, 0.945] | -0.005 [-0.012, -0.003] | 0.030 |
| AP trunk acc. SampEn | -0.007 [-0.019, 0.007] | -0.068 [-0.240, 0.117] | 0.110 [0.103, 0.118] | 0.177 |
| ML trunk acc. SampEn | 0.119 [0.093, 0.211] | 0.290 [0.180, 0.430] | -0.082 [-0.099, -0.066] | 0.186 |
| VT trunk acc. SampEn | 0.341 [0.098, 0.527] | 0.353 [0.107, 0.632] | -0.010 [-0.016, 0.025] | 0.109 |

**Notes:** See **Table 1**. $\lambda_{\text{short}}$ : maximum Lyapunov exponent. ACI: attractor complexity index. SampEn: sample entropy.

ACI showed the best agreement among trunk-level nonlinear metrics. Vertical and anteroposterior ACI reached good agreement (ICC = 0.843 and 0.818) with small negative biases and high correlations (r ≥ 0.899). Mediolateral ACI was slightly lower (ICC = 0.750) but remained in the good range.

λshort agreement depended strongly on the acceleration direction. Anteroposterior trunk λshort showed moderate agreement (median ICC = 0.639), with small positive biases and moderate correlations (r = 0.641). Vertical trunk λshort showed poor to moderate agreement (median ICC = 0.383) despite high correlations (median r = 0.797), reflecting a substantial systematic positive bias (median = 0.184) that degraded absolute agreement. Mediolateral trunk λshort showed poor agreement (median ICC = 0.083), with a large systematic positive bias (median = 0.371) that dominated inter-system discrepancy.

SampEn showed poor agreement for all three trunk acceleration directions. Anteroposterior SampEn showed near-zero or negative ICCs (median = −0.007) with a large positive bias (median = 0.110), indicating that the markerless system produced systematically higher entropy values with no preservation of inter-individual ranking. Mediolateral SampEn (median ICC = 0.119) and vertical SampEn (median ICC = 0.341) followed the same pattern, with biases in opposing directions.

### 3.3. Effect of walking speed on inter-system agreement

Of the 45 gait measures tested, 19 showed a significant effect of walking speed on inter-system bias after FDR correction (pFDR < 0.05, **Supplementary Table S9**). Among linear measures, step speed and step length biases increased progressively with belt speed (η²p = 0.63 and 0.50), with the self-paced condition not differing from 100% PWS. Step width showed a smaller positive bias at 140% PWS (η²p = 0.18). Among variability metrics, double-support time SD was the only measure significantly affected (η²p = 0.32), with a larger bias in the self-paced condition. Among DFA exponents, only body position α reached significance (η²p = 0.22), with a larger negative bias in the self-paced condition.

At the joint level, ankle λshort showed a reduction in negative bias magnitude at faster speeds for both methods (η²p = 0.40 and 0.25). Smaller effects (η²p = 0.18 - 0.32) were also found for hip SampEn, hip λshort (Pose2Sim), knee SampEn (both methods), and hip ACI (both methods).

Among trunk nonlinear measures, mediolateral λshort showed the largest speed effect (η²p = 0.60), with positive bias increasing from 0.35 at 100% PWS to 0.47 at 140% PWS. Anteroposterior λshort followed a similar pattern (η²p = 0.37). Smaller effects were observed for vertical ACI (η²p = 0.31), mediolateral ACI (η²p = 0.19), mediolateral SampEn (η²p = 0.21), and vertical SampEn (η²p = 0.20).

## DISCUSSION

This study evaluated the validity of a multi-view markerless system against a marker-based reference for (i) linear measures of conventional gait variables, namely spatiotemporal parameters and sagittal-plane joint kinematics, and (ii) nonlinear measures indexing the temporal structure of spatiotemporal parameters and of joint angle and trunk acceleration time series – the latter not previously validated for a video-based markerless system. Validity was not uniform but depended on the measure and followed a tiered pattern. The spatiotemporal parameters agreed closely enough to replace the marker-based values, apart from a small systematic offset in the spatial ones. The sagittal-plane angles were also valid, with the error increasing from the hip to the ankle. Among the nonlinear metrics, the Attractor Complexity Index (ACI) agreed well for every continuous signal, as did the DFA exponents of most step-based series, whereas the maximum Lyapunov exponent and sample entropy (SampEn) differed in absolute value between systems even though they still ranked participants consistently. Validity is therefore measure-specific, and each measure must be assessed on its own rather than inferred from the overall performance of the system.

Two factors explain this tiered pattern. The first is the time scale that each measure examines. The second is the size of the true signal relative to the measurement error added by the markerless system. Because this error is roughly constant and does not grow with the signal, it matters little when the signal is large but dominates when the signal is small. ACI is computed over a window of 4 to 10 strides (Terrier, 2019; Terrier & Reynard, 2018) that spans the whole gait cycle, so it is barely affected by small and rapid fluctuations. It therefore remained valid at every joint, in every trunk direction, and for trunk position, velocity, and acceleration (**Supplementary Tables S6 to S8**). The DFA exponents share this advantage, because they are computed across many steps rather than from fine detail. Most of them agreed well between systems, and the markerless system lowered them only slightly, as expected when frame-to-frame noise weakens long-range correlation (Damouras et al., 2010; Phinyomark et al., 2020). In contrast, λshort and SampEn depend on the fine detail of the signal, which noise, sampling, and filtering distort (Mehdizadeh & Sanjari, 2017; Raffalt et al., 2020; Yentes & Raffalt, 2021). At the hip and knee they still ranked participants correctly (Pearson’s r above ICC) but were shifted by a constant bias. For the trunk, they deteriorated with each successive derivative, because differentiation amplifies high-frequency noise (Winter, 2009). SampEn shows this most clearly, agreeing almost perfectly for trunk position but poorly for trunk acceleration (**Supplementary Tables S6 to S8**). A small signal amplitude has the same effect. Because mediolateral motion is limited during treadmill walking, the reconstruction error forms a large share of the mediolateral signal, and these measures were accordingly the least reliable. This was true of step width and its variability, of the mediolateral component of body-position α, which the markerless system underestimated, and of mediolateral trunk λshort, whose bias grew as faster belts increased mediolateral motion. For the conventional gait variables, the present results match or exceed those of earlier markerless studies.

The spatiotemporal parameters were almost identical because gait events were detected the same way in both systems. The small positive bias in the spatial parameters matches the known offset between skin markers and pose-estimated keypoints (Kanko, Laende, Strutzenberger, et al., 2021; Stenum et al., 2021). This bias left the ranking of participants unchanged but grew with belt speed, so absolute spatial comparisons across speeds should be corrected for it. At 140% of preferred speed, the step-speed MDCmax already approached a clinically meaningful gait-speed difference (Bohannon & Glenney, 2014; Perera et al., 2006). The sagittal-plane angle errors remained close to the usual 5° limit at every joint (McGinley et al., 2009), reaching it only at the ankle with Pose2Sim (5.1°). Agreement was best at the hip, with knee and ankle showing comparable, higher error than hip, as observed in earlier work (Drazan et al., 2021; Kanko, Laende, Davis, et al., 2021). The hip showed no difference between systems, while the two angle methods produced opposite bias directions at the ankle.

These results also clarify where markerless capture is most useful. Because it needs no physical markers, it avoids the placement errors that vary between operators and sessions (Gorton et al., 2009), and is faster and less expensive to set up. It also supports the long, many-step recordings that nonlinear analysis requires. Together, these properties make it well suited to repeated, clinical, and field measurements (Uhlrich et al., 2023; Wade et al., 2022). Its accuracy is nonetheless limited at two stages. Pose-estimation depends on occlusion, lighting, and the 2D model, and it locates keypoints on a fixed skeleton rather than on palpated anatomical landmarks, so a residual localization error is unavoidable (Needham, Evans, Cosker, & Colyer, 2021; Needham, Evans, Cosker, Wade, et al., 2021). An optional model-fitting step such as Pose2Sim can smooth the trajectories, but it reintroduces the modeling assumptions that a markerless approach is meant to avoid (Delp et al., 2007; Pagnon et al., 2022a). The model-free method used here therefore keeps the pipeline light while matching Pose2Sim for most measures.

This study has some limitations. The participants were healthy young adults, and the ranking of the noise-sensitive measures may differ in slower, more variable, or pathological gait, where fine-scale dynamics carry the most clinical information. All data were recorded on a treadmill, which standardizes acquisition but constrains the DFA (Damouras et al., 2010) and limits how far the results apply to overground walking. The three-camera configuration and the fixed pose-estimation model, although chosen to reflect current best practice in multi-view markerless capture, represent one specific implementation, and different camera placements, numbers, or 2D pose models will modify the validity profile (D’Souza et al., 2024; Stenum et al., 2021). Finally, validation was limited to the sagittal plane, because the frontal-and transverse-plane angles of the Plug-in-Gait model are themselves affected by soft-tissue and cross-talk errors and cannot serve as a clean reference (Chafetz et al., 2026; Peters et al., 2010).

## CONCLUSION

This markerless system supports interchangeable measurement of temporal parameters, group-level estimation of most spatial parameters with a speed-dependent offset, and valid sagittal-plane hip and knee kinematics, whereas ankle kinematics showed opposite bias directions between methods despite comparable RMSE magnitude. Among nonlinear metrics, ACI is valid across all joints and trunk directions, and is therefore the most reliable nonlinear measure to compare with marker-based data or across systems. DFA scaling exponents for step time, step length, step speed, and body position support group-level comparisons, provided that the systematic attenuation of absolute values is acknowledged. At the hip and knee, λshort and SampEn support between-group comparison within the markerless system but are not interchangeable with marker-based values. Finally, agreement was weakest for ankle λshort, ankle SampEn, and trunk SampEn, which are not yet reliable with the current setup. This limitation is technical, however, and denser camera coverage or refined preprocessing could bring them within reach. Together, these results give markerless gait analysis a tiered, measure-specific scope that reaches well beyond spatiotemporal description into the nonlinear dynamics of gait.

## ACKNOWLEDGMENTS

The authors gratefully thank all study participants for their time, commitment, and invaluable contribution to this research.

The authors also acknowledge the COMETE laboratory (UMR-S 1075, INSERM / Université de Caen Normandie), the CIREVE (Interdisciplinary Centre for Virtual Reality), and the company a-gO for their essential logistical, administrative, and technical support in facilitating the implementation of the study.

The authors also warmly thank Sophie Madeleine, Director of CIREVE, and Philippe Fleury, founder and former Director of CIREVE, whose vision and support made possible the joint acquisition of the M-Gait instrumented dual-belt treadmill and the motorized fall-arrest system used in this study, funded through the TRISIM project (CPER 2021-2027).

## AUTHOR CONTRIBUTIONS

**Baptiste Perthuy:** Conceptualization, Data curation, Formal analysis, Investigation, Methodology, Project administration, Software, Visualization, Writing – original draft. **Hugues Vinzant:** Conceptualization, Methodology, Resources, Software, Supervision, Writing – review & editing. **Clément Brifault:** Conceptualization, Methodology, Resources, Software, Writing – review & editing. **Nicolas Lefèvre:** Methodology, Resources, Writing – review & editing. **Alexandre Dalibot:** Conceptualization, Methodology, Resources, Supervision, Writing – review & editing. **Sofiane Ramdani:** Methodology, Supervision, Validation, Writing – review & editing. **Leslie M. Decker:** Conceptualization, Funding acquisition, Methodology, Project administration, Resources, Supervision, Validation, Writing – review & editing.

## STATEMENTS AND DECLARATIONS

No conflicts of interest, financial or otherwise, are declared by the authors.

## DATA AVAILABILITY STATEMENT

The data that support the findings of this study are available within the article and its Supplementary Materials.

