## Supplementary Materials for "From linear to nonlinear gait measures: validity of a multi-view markerless motion capture system"

**Appendix A:** Nonlinear analyses

*Detrended fluctuation analysis*

Long-range temporal correlations in step-to-step fluctuations were quantified using Detrended Fluctuation Analysis (DFA; Peng et al., 1995). DFA was applied to step-based time series, namely step time, step length, step speed, step width, and body position. The number of consecutive steps analyzed ($N$) was kept identical across systems, while varying across participants and walking conditions (range 512 - 1024 steps).

Given a step-based series $x\left( i \right), i=1, \ldots, N$, with mean $\bar{x}$, the signal was first integrated to obtain cumulative profile,

$$y\left( k \right)=\sum_{i=1}^{k} \left[ x\left( i \right)-\bar{x} \right], k=1, \ldots, N$$

The profile $y(k)$ was then divided into non-overlapping windows of length $n$. Within each window a local polynomial trend $y_{n}(k)$ was fitted by ordinary least squares and subtracted, and the fluctuation function was computed as the root-mean-square of the detrended profile,

$$F\left( n \right)=\sqrt{\frac{1}{N}\sum_{k=1}^{N} \left[ y\left( k \right)-y_{n}(k) \right]^{2}}$$

This procedure was repeated across window sizes ranging from 16 steps to $N/9$, sampled in even (arithmetic) increments of six steps, following the evenly-spaced recommendations for gait time series (Damouras et al., 2010; Phinyomark et al., 2020). Because $F(n)$ scales with window size as a power law, $F\left( n \right) \sim n^{\alpha}$, the scaling exponent α was estimated as the slope of the linear regression of $\log F(n)$on $\log n$. Values $\alpha> 0.5$ indicate persistent correlations, whereby fluctuations tend to be followed by changes in the same direction, whereas $\alpha< 0.5$ reflects antipersistent behavior, with successive changes more likely to alternate in sign. Values $\alpha\approx0.5$ corresponds to uncorrelated, white-noise-like fluctuations (Hausdorff et al., 1996; Peng et al., 1995).

The detrending order determines which polynomial trends are removed within each window: DFA-l eliminates trends up to order l in the integrated profile (equivalently, up to order l-1 in the original series). Although first-order (linear, DFA-1) detrending is the most common default in stride-interval studies, second-order detrending (DFA-2) was adopted here in accordance with the parameter recommendations of Phinyomark et al. (2020) for time series of the length analyzed (512 - 1024 steps). The same detrending order was applied identically across systems, participants and conditions, so that between-system comparisons remained internally consistent.

*Sample entropy*

Signal complexity was quantified using Sample Entropy (SampEn, Richman & Moorman, 2000), defined as the negative logarithm of the conditional probability that two sequences of length $m$ that match within a tolerance $r$ remain similar when extended to $m+1$ points. SampEn was computed on the continuous trunk-acceleration signals and on the sagittal-plane hip, knee and ankle joint angles.

For a time series $u(1), \ldots, u(N)$, template vectors were formed as $X_{m}(i)=[u(i), u(i+1), \ldots, u(i+m-1)]$, for $i=1, \ldots, N-m+1$. For each pair $i\neq j$, similarity was evaluated using the Chebyshev distance,

$$d\left[ X_{m}\left( i \right), X_{m}\left( j \right) \right]=\max_{k=0, \ldots, m-1} |u(i+k)-u(j+k)|\leq r$$

Letting $B^{m}(r)$ and $A^{m}(r)$ denote the average numbers of within-tolerance matches of length $m$ and $m+1$ respectively (self-matched excluded), sample entropy is,

$$SampEn\left( m, r, N \right)=-\ln\left[ \frac{A^{m}\left( r \right)}{B^{m}\left( r \right)} \right]$$

The embedding dimension was fixed at $m=2$ and the tolerance at $r=0.2\times SD$ of the individual time series, in line with parameter choices validated for human gait signals (Ahmadi et al., 2018; McCamley et al., 2018; Yentes et al., 2013). To confirm that these values were appropriate, SampEn was recomputed across a grid of $m\in[2-5]$ and $r\in[0.10-0.50] \times SD$ values and inspected visually; the selected parameters fell within the plateau region in which entropy estimates are stable, as typically reported for gait data. Lower SampEn values indicate greater regularity and predictability, whereas higher values reflect greater irregularity and complexity.

*Maximum Lyapunov exponent*

Divergence of movement trajectories was assessed from the continuous trunk-acceleration signals and on the sagittal-plane hip, knee and ankle joint angles. Each scalar signal $s(1), \ldots, s(N)$ was first reconstructed in state space using time-delay embedding (Abarbanel, 1996; Kantz & Schreiber, 2003; Takens, 1981). State vectors were built as,

$$X\left( i \right)=[s\left( i \right), s\left( i+\tau\right), s\left( 1+2\tau\right), \ldots, s\left( i+\left( m-1 \right)\tau\right]$$

Where $\tau$ is the time delay and $m$ the embedding dimension. By Takens’ theorem, for a suitable delay and sufficiently large embedding dimension, the reconstructed attractor is topologically equivalent to that of the original system and preserves its invariant dynamical properties, including divergence rates.

The time delay $\tau$ was set as the first local minimum of the average mutual information (AMI) function (Fraser & Swinney, 1986). Unlike the linear autocorrelation, which only captures second-order (linear) dependence, AMI is computed from the joint probability distribution of $s\left( i \right)$ and its delayed value $s\left( i+\tau\right)$ and therefore quantifies their full statistical dependence, including nonlinear, higher-order relationships (Abarbanel, 1996; Kantz & Schreiber, 2003). Using the first local minimum selects the delay at which $s\left( i+\tau\right)$ adds the newest information while remaining dynamically related to $s\left( i \right)$.

The embedding dimension $m$ was set to the smallest value for which the proportion of false nearest neighbors fell below 5% (Kennel et al., 1992); a neighbor is deemed false when increasing the dimension from $m$ to $m+1$ separates it beyond a threshold, indicating that its proximity was an artefact of projecting the attractor onto too few dimensions.

For each signal (each acceleration axis and joint angle) and each walking condition, a single time delay and embedding dimension were used, taken as the median of the values estimated across all participants. These common parameters were applied to both systems and to all participants, so that any difference in λshort would reflect the underlying dynamics rather than differences in state-space reconstruction, while still being allowed to vary across signals and walking conditions (Dingwell & Cusumano, 2000; Raffalt et al., 2019). Each time series was time-normalized to 101 samples per stride to ensure a consistent temporal resolution across participants and to minimize the influence of stride-to-stride duration variability (Raffalt et al., 2019).

Trajectory divergence was quantified using the short-term maximum Lyapunov exponent (λshort) with the algorithm of Rosenstein et al. (1993). For each reference vector $X(i)$, the nearest neighbor $X(\hat{j})$ was identified by Euclidean distance, excluding temporally adjacent points through a Theiler window equal to one stride. The Euclidean distance between each pair of initially neighboring trajectories, $d_{i}\left( k \right)=\left\| X\left( i+k \right)-X(\hat{j}+k) \right\|$, was tracked over $k$ discrete steps, and the mean logarithmic divergence curve was obtained by averaging the natural logarithm across all $M$ reference vectors, where $\left\langle\cdot\right\rangle$ denotes the average over all reference vectors $i=1, \ldots, M$,

$$\left\langle\ln d(k) \right\rangle=\frac{1}{M}\sum_{i=1}^{M} \ln d_{i}(k)$$

λshort was estimated as the slope of the linear region of this curve over the first 0 - 0.5 strides cycles. Higher λshort values reflect faster local divergence of neighboring trajectories, and thus lower local dynamic stability, whereas smaller values indicate more stable locomotor dynamics.

*Attractor complexity index*

Long-term divergence characteristics were finally quantified using the Attractor Complexity Index (ACI) (Terrier & Reynard, 2018), derived from the same divergence curves. Whereas λshort was computed over the initial 0 - 0.5 strides, the ACI was defined as the slope of the mean logarithmic divergence curve within the 4 - 10 stride intervals, thereby capturing the slower divergence dynamics of the attractor. Higher ACI values indicate a more complex gait dynamic with wider attractor boundaries that permit greater long-term divergence, whereas lower values reflect a more restricted attractor and reduced complexity of gait control (Terrier, 2019; Terrier & Reynard, 2018).

**Supplementary Table S1.** Step-based spatiotemporal parameters, definitions and computation formulas

| **Parameters** | **Definition** | **Computation formula** |
| --- | --- | --- |
| Step time (s) | Time interval between two consecutive heel strikes of opposite feet | $Step {time}_{n}=t_{HS}^{(n+1)}-t_{HS}^{(n)}$ |
| Step length (m) | Anteroposterior distance between the heel markers of opposite feet measured at heel strike | $Step {length}_{n}=\left\vert{Heel}_{AP}^{(n+1)}-{Heel}_{AP}^{(n)} \right\vert$ |
| Step speed (m/s) | Forward velocity of the body during a given step | $Step {speed}_{n}=\frac{Step {length}_{n}}{Step {time}_{n}}$ |
| Step width (m) | Mediolateral distance between the heel markers of opposite feet measured at heel strike | $Step {width}_{n}=\left\vert{Heel}_{ML}^{(n+1)}-{Heel}_{ML}^{(n)} \right\vert$ |
| Body position (m) | Mediolateral position of the body at each step, defined as the midpoint between the left and right heel markers | $Body {position}_{n}=0.5\times({Heel}_{ML}^{(n)}+{Heel}_{ML}^{(n+1)})$ |
| Single-support time (s) | Time period during which only one foot is in contact with the ground | $Single {support time}_{n}=t_{HS}^{(n)}-t_{TO}^{(n-1)}$ |
| Double-support time (s) | Time period during which both feet are simultaneously in contact with the ground | $Double {support time}_{n}={IDS}_{n}+{TDS}_{n}$ |
| Cadence (step/min) | Number of steps performed per minute | $Cadence=60 / \bar{Step time}$ |

**Notes:** $t_{HS}^{(n)}$ and $t_{TO}^{(n)}$ denote the timestamps of the $n^{th}$ heel strike and toe-off events, respectively. ${Heel}_{AP/ML}^{(n)}$ denote the anteroposterior (AP) or mediolateral (ML) position of the heel marker at the $n^{th}$ heel-strike events.

**Supplementary Table S2.** Agreement between marker-based and markerless systems for mean of spatiotemporal parameters across walking conditions.

| **Measure** | **Speed** | **Vicon**  **(Mean ± SD)** | **Markerless (Mean ± SD)** | | **Bias (95% LoA)** | | **Prop. bias**  **(slope, p)** | **ICC (A, 1)** | | **r** | **SEM** | **MDC** |
| --- | --- | --- | --- | --- | --- | --- | --- | --- | --- | --- | --- | --- |
| Step speed (m/s) | 100 % PWS | 1.154 ± 0.074 | 1.174 ± 0.077 | | 0.021 (-0.006, 0.048) | | 0.033, p = 0.418 | 0.949 [0.844, 0.978] | | 0.984 | 0.017 | 0.047 |
|  | 120 % PWS | 1.373 ± 0.088 | 1.401 ± 0.091 | | 0.028 (0.001, 0.055) | | 0.035, p = 0.316 | 0.941 [0.825, 0.974] | | 0.988 | 0.022 | 0.060 |
|  | 140 % PWS | 1.586 ± 0.100 | 1.623 ± 0.106 | | 0.037 (0.005, 0.070) | | 0.060, p = 0.086 | 0.927 [0.787, 0.967] | | 0.989 | 0.028 | 0.078 |
|  | Self-paced | 1.233 ± 0.173 | 1.257 ± 0.183 | | 0.025 (-0.006, 0.055) | | 0.055, **p = 0.002** | 0.987 [0.960, 0.994] | | 0.998 | 0.020 | 0.056 |
| Step time (s) | 100 % PWS | 0.569 ± 0.038 | 0.569 ± 0.038 | | 0.000 (-0.000, 0.000) | | -0.000, p = 0.209 | 1.000 [1.000, 1.000] | | 1.000 | 0.000 | 0.000 |
|  | 120 % PWS | 0.526 ± 0.036 | 0.526 ± 0.036 | | 0.000 (0.000, 0.000) | | 0.000, p = 0.208 | 1.000 [1.000, 1.000] | | 1.000 | 0.000 | 0.000 |
|  | 140 % PWS | 0.491 ± 0.035 | 0.491 ± 0.035 | | 0.000 (-0.000, 0.000) | | 0.000, p = 0.285 | 1.000 [1.000, 1.000] | | 1.000 | 0.000 | 0.000 |
|  | Self-paced | 0.553 ± 0.049 | 0.553 ± 0.049 | | 0.000 (-0.000, 0.000) | | 0.000, p = 0.507 | 1.000 [1.000, 1.000] | | 1.000 | 0.000 | 0.000 |
| Step length (m) | 100 % PWS | 0.655 ± 0.046 | 0.666 ± 0.048 | | 0.012 (-0.004, 0.027) | | 0.035, p = 0.353 | 0.957 [0.889, 0.977] | | 0.986 | 0.010 | 0.027 |
|  | 120 % PWS | 0.721 ± 0.050 | 0.736 ± 0.052 | | 0.015 (0.000, 0.029) | | 0.026, p = 0.434 | 0.950 [0.872, 0.971] | | 0.990 | 0.011 | 0.032 |
|  | 140 % PWS | 0.777 ± 0.054 | 0.795 ± 0.057 | | 0.018 (0.003, 0.034) | | 0.048, p = 0.129 | 0.939 [0.856, 0.964] | | 0.991 | 0.014 | 0.038 |
|  | Self-paced | 0.674 ± 0.068 | 0.687 ± 0.073 | | 0.013 (-0.003, 0.029) | | 0.073, **p = 0.003** | 0.976 [0.941, 0.988] | | 0.996 | 0.011 | 0.030 |
| Step width (m) | 100 % PWS | 0.107 ± 0.026 | 0.114 ± 0.025 | | 0.007 (-0.012, 0.026) | | -0.043, p = 0.631 | 0.893 [0.756, 0.952] | | 0.927 | 0.008 | 0.023 |
|  | 120 % PWS | 0.106 ± 0.022 | 0.112 ± 0.022 | | 0.006 (-0.012, 0.025) | | -0.011, p = 0.916 | 0.873 [0.700, 0.947] | | 0.906 | 0.008 | 0.022 |
|  | 140 % PWS | 0.108 ± 0.021 | 0.114 ± 0.021 | | 0.006 (-0.013, 0.024) | | 0.011, p = 0.918 | 0.876 [0.703, 0.949] | | 0.902 | 0.007 | 0.021 |
|  | Self-paced | 0.111 ± 0.024 | 0.117 ± 0.023 | | 0.007 (-0.012, 0.025) | | -0.025, p = 0.792 | 0.887 [0.745, 0.953] | | 0.919 | 0.008 | 0.022 |
| Double-support time (s) | 100 % PWS | 0.367 ± 0.040 | 0.369 ± 0.039 | | 0.002 (-0.029, 0.034) | | 0.040, p = 0.675 | 0.915 [0.801, 0.966] | | 0.914 | 0.011 | 0.031 |
|  | 120 % PWS | 0.313 ± 0.033 | 0.312 ± 0.033 | | -0.000 (-0.025, 0.024) | | 0.010, p = 0.901 | 0.933 [0.850, 0.969] | | 0.930 | 0.008 | 0.023 |
|  | 140 % PWS | 0.271 ± 0.030 | 0.272 ± 0.032 | | 0.000 (-0.016, 0.017) | | 0.062, p = 0.318 | 0.936 [0.932, 0.979] | | 0.964 | 0.006 | 0.016 |
|  | Self-paced | 0.343 ± 0.058 | 0.346 ± 0.061 | | 0.002 (-0.025, 0.030) | | 0.043, p = 0.442 | 0.972 [0.932, 0.988] | | 0.972 | 0.009 | 0.027 |
| Single-support time (s) | 100 % PWS | 0.385 ± 0.023 | 0.384 ± 0.023 | | -0.001 (-0.017, 0.015) | | -0.028, p = 0.723 | 0.941 [0.796, 0.979] | | 0.940 | 0.006 | 0.015 |
|  | 120 % PWS | 0.370 ± 0.022 | 0.370 ± 0.022 | | 0.000 (-0.012, 0.012) | | 0.010, p = 0.878 | 0.964 [0.890, 0.987] | | 0.962 | 0.004 | 0.012 |
|  | 140 % PWS | 0.355 ± 0.022 | 0.355 ± 0.021 | | -0.000 (-0.009, 0.008) | | -0.064, p = 0.151 | 0.981 [0.937, 0.992] | | 0.982 | 0.003 | 0.008 |
|  | Self-paced | 0.381 ± 0.025 | 0.380 ± 0.023 | | -0.001 (-0.015, 0.013) | | -0.084, p = 0.220 | 0.956 [0.892, 0.985] | | 0.959 | 0.005 | 0.014 |
| Cadence (steps/min) | 100 % PWS | 105.99 ± 6.90 | 105.98 ± 6.90 | | -0.004 (-0.011, 0.004) | | -0.000, p = 0.301 | 1.000 [1.000, 1.000] | | 1.000 | 0.004 | 0.010 |
|  | 120 % PWS | 114.46 ± 7.40 | 114.46 ± 7.40 | | -0.003 (-0.005, -0.001) | | -0.000, p = 0.464 | 1.000 [1.000, 1.000] | | 1.000 | 0.002 | 0.006 |
|  | 140 % PWS | 122.75 ± 8.31 | 122.74 ± 8.31 | | -0.003 (-0.008, 0.003) | | 0.000, p = 0.453 | 1.000 [1.000, 1.000] | | 1.000 | 0.003 | 0.007 |
|  | Self-paced | 109.41 ± 9.22 | 109.41 ± 9.22 | | -0.000 (-0.011, 0.011) | | 0.000, p = 0.895 | 1.000 [1.000, 1.000] | | 1.000 | 0.004 | 0.011 |
| ICC (A, 1) | < 0.50, poor agreement | | | 0.50 – 0.75, moderate agreement | | 0.75 – 0.90, good agreement | | | > 0.90 excellent agreement | | | |

**Notes:** Values are group means ± SD. Bias and 95% limits of agreement (LoA) were derived from Bland-Altman analysis. Proportional bias p-values reflect regression of inter-system difference on the measurement mean; bold indicates p < 0.05. ICC (A, 1): intraclass correlation coefficient, two-way mixed-effects model, absolute agreement, single measure. SEM: standard error of measurement. MDC: minimal detectable change. PWS: preferred walking speed.

**Supplementary Table S3.** Agreement between marker-based and markerless systems for standard deviation of spatiotemporal parameters across walking conditions.

| **Measure** | **Speed** | **Vicon**  **(Mean ± SD)** | **Markerless (Mean ± SD)** | | **Bias (95% LoA)** | | **Prop. bias**  **(slope, p)** | **ICC (A, 1)** | | **r** | | **SEM** | **MDC** |
| --- | --- | --- | --- | --- | --- | --- | --- | --- | --- | --- | --- | --- | --- |
| Step speed variability (m/s) | 100PWS | 0.033 ± 0.008 | 0.036 ± 0.011 | | 0.003 (-0.006, 0.012) | | 0.250, **p = 0.025** | 0.836 [0.676, 0.923] | | 0.896 | | 0.004 | 0.011 |
|  | 120PWS | 0.034 ± 0.008 | 0.038 ± 0.010 | | 0.004 (-0.006, 0.014) | | 0.259, **p = 0.040** | 0.756 [0.575, 0.877] | | 0.865 | | 0.004 | 0.012 |
|  | 140PWS | 0.038 ± 0.010 | 0.041 ± 0.013 | | 0.004 (-0.008, 0.015) | | 0.303, **p = 0.003** | 0.840 [0.768, 0.923] | | 0.917 | | 0.005 | 0.013 |
|  | Self-paced | 0.070 ± 0.028 | 0.073 ± 0.029 | | 0.003 (-0.004, 0.009) | | 0.031, p = 0.244 | 0.989 [0.947, 0.996] | | 0.994 | | 0.003 | 0.008 |
| Step time variability (s) | 100PWS | 0.012 ± 0.003 | 0.012 ± 0.003 | | 0.000 (-0.002, 0.003) | | 0.061, p = 0.562 | 0.898 [0.744, 0.976] | | 0.898 | | 0.001 | 0.003 |
|  | 120PWS | 0.009 ± 0.002 | 0.010 ± 0.002 | | 0.001 (-0.002, 0.003) | | 0.349, **p = 0.006** | 0.804 [0.625, 0.944] | | 0.869 | | 0.001 | 0.003 |
|  | 140PWS | 0.009 ± 0.004 | 0.009 ± 0.004 | | 0.000 (-0.001, 0.002) | | 0.015, p = 0.702 | 0.983 [0.753, 0.997] | | 0.986 | | 0.001 | 0.002 |
|  | Self-paced | 0.015 ± 0.007 | 0.015 ± 0.006 | | 0.000 (-0.001, 0.001) | | -0.004, p = 0.869 | 0.995 [0.968, 0.999] | | 0.995 | | 0.000 | 0.001 |
| Step length variability (m) | 100PWS | 0.015 ± 0.006 | 0.017 ± 0.006 | | 0.002 (-0.002, 0.005) | | 0.067, p = 0.362 | 0.918 [0.702, 0.974] | | 0.950 | | 0.002 | 0.005 |
|  | 120PWS | 0.015 ± 0.005 | 0.016 ± 0.005 | | 0.002 (-0.002, 0.005) | | 0.158, p = 0.062 | 0.879 [0.760, 0.952] | | 0.937 | | 0.002 | 0.005 |
|  | 140PWS | 0.015 ± 0.004 | 0.016 ± 0.005 | | 0.002 (-0.002, 0.006) | | 0.114, p = 0.248 | 0.863 [0.762, 0.933] | | 0.911 | | 0.002 | 0.005 |
|  | Self-paced | 0.028 ± 0.010 | 0.029 ± 0.010 | | 0.002 (-0.001, 0.004) | | 0.020, p = 0.565 | 0.978 [0.890, 0.992] | | 0.989 | | 0.001 | 0.004 |
| Step width variability (m) | 100PWS | 0.021 ± 0.004 | 0.022 ± 0.004 | | 0.001 (-0.002, 0.004) | | -0.017, p = 0.844 | 0.916 [0.762, 0.980] | | 0.928 | | 0.001 | 0.004 |
|  | 120PWS | 0.021 ± 0.004 | 0.023 ± 0.005 | | 0.002 (-0.004, 0.008) | | 0.362, **p = 0.039** | 0.643 [0.302, 0.865] | | 0.751 | | 0.003 | 0.007 |
|  | 140PWS | 0.022 ± 0.003 | 0.024 ± 0.004 | | 0.002 (-0.005, 0.009) | | 0.155, p = 0.518 | 0.489 [0.028, 0.849] | | 0.560 | | 0.003 | 0.008 |
|  | Self-paced | 0.021 ± 0.005 | 0.022 ± 0.005 | | 0.001 (-0.004, 0.006) | | 0.014, p = 0.912 | 0.851 [0.560, 0.980] | | 0.858 | | 0.002 | 0.005 |
| Double-support time variability (s) | 100PWS | 0.013 ± 0.003 | 0.013 ± 0.003 | | 0.000 (-0.001, 0.002) | | 0.020, p = 0.737 | 0.964 [0.921 0.985] | | 0.970 | | 0.000 | 0.002 |
|  | 120PWS | 0.010 ± 0.002 | 0.011 ± 0.002 | | 0.000 (-0.000, 0.001) | | 0.015, **p = 0.015** | 0.954 [0.904, 0.983] | | 0.969 | | 0.000 | 0.001 |
|  | 140PWS | 0.010 ± 0.002 | 0.010 ± 0.002 | | 0.000 (-0.000, 0.000) | | -0.000, p = 0.999 | 0.989 [0.968, 0.997] | | 0.991 | | 0.000 | 0.000 |
|  | Self-paced | 0.020 ± 0.010 | 0.020 ± 0.010 | | 0.000 (-0.001, 0.002) | | 0.038, **p = 0.044** | 0.991 [0.959, 0.997] | | 0.996 | | 0.000 | 0.002 |
| Single-support time variability (s) | 100PWS | 0.008 ± 0.002 | 0.008 ± 0.002 | | 0.001 (-0.002, 0.003) | | 0.319, **p = 0.048** | 0.723 [0.428, 0.964] | | 0.786 | | 0.001 | 0.003 |
|  | 120PWS | 0.007 ± 0.001 | 0.007 ± 0.001 | | 0.000 (-0.001, 0.001) | | 0.088, p = 0.363 | 0.900 [0.725, 0.963] | | 0.914 | | 0.000 | 0.001 |
|  | 140PWS | 0.007 ± 0.005 | 0.007 ± 0.005 | | 0.000 (-0.001, 0.001) | | 0.003, p = 0.863 | 0.997 [0.816, 0.999] | | 0.997 | | 0.000 | 0.001 |
|  | Self-paced | 0.009 ± 0.002 | 0.009 ± 0.002 | | 0.000 (-0.001, 0.001) | | 0.041, p = 0.597 | 0.947 [0.834, 0.984] | | 0.945 | | 0.000 | 0.001 |
| ICC (A, 1) | < 0.50, poor agreement | | | 0.50 – 0.75, moderate agreement | | 0.75 – 0.90, good agreement | | | > 0.90 excellent agreement | | | | |

**Notes:** Values are group means ± SD. Bias and 95% limits of agreement (LoA) were derived from Bland-Altman analysis. Proportional bias p-values reflect regression of inter-system difference on the measurement mean; bold indicates p < 0.05. ICC (A, 1): intraclass correlation coefficient, two-way mixed-effects model, absolute agreement, single measure. SEM: standard error of measurement. MDC: minimal detectable change. PWS: preferred walking speed.

**Supplementary Table S4.** Agreement between marker-based and markerless systems for temporal structure of spatiotemporal parameters across walking conditions.

| **Measure** | **Speed** | **Vicon**  **(Mean ± SD)** | **Markerless (Mean ± SD)** | | **Bias (95% LoA)** | | **Prop. bias**  **(slope, p)** | | **ICC (A, 1)** | | | **r** | | **SEM** | | **MDC** |
| --- | --- | --- | --- | --- | --- | --- | --- | --- | --- | --- | --- | --- | --- | --- | --- | --- |
| Step speed α | 100PWS | 0.585 ± 0.126 | 0.577 ± 0.144 | | -0.008 (-0.181, 0.165) | | 0.150, p = 0.330 | | 0.793 [0.541, 0.920] | | | 0.794 | | 0.061 | | 0.169 |
|  | 120PWS | 0.573 ± 0.114 | 0.551 ± 0.132 | | -0.022 (-0.148, 0.104) | | 0.154, p = 0.200 | | 0.856 [0.750, 0.932] | | | 0.873 | | 0.046 | | 0.129 |
|  | 140PWS | 0.587 ± 0.107 | 0.582 ± 0.106 | | -0.006 (-0.111, 0.100) | | -0.010, p = 0.932 | | 0.876 [0.792, 0.918] | | | 0.872 | | 0.037 | | 0.103 |
|  | Self-paced | 1.001 ± 0.165 | 0.986 ± 0.160 | | -0.015 (-0.157, 0.127) | | -0.032, p = 0.764 | | 0.902 [0.735, 0.969] | | | 0.902 | | 0.050 | | 0.140 |
| Step time α | 100PWS | 0.597 ± 0.108 | 0.589 ± 0.118 | | -0.008 (-0.117, 0.101) | | 0.094, p = 0.411 | | 0.882 [0.733, 0.955] | | | 0.882 | | 0.038 | | 0.106 |
|  | 120PWS | 0.574 ± 0.057 | 0.548 ± 0.086 | | -0.026 (-0.135, 0.083) | | 0.463, **p = 0.007** | | 0.676 [0.433, 0.852] | | | 0.772 | | 0.042 | | 0.116 |
|  | 140PWS | 0.584 ± 0.105 | 0.569 ± 0.116 | | -0.015 (-0.100, 0.069) | | 0.103, p = 0.244 | | 0.919 [0.811, 0.974] | | | 0.929 | | 0.031 | | 0.087 |
|  | Self-paced | 0.772 ± 0.131 | 0.764 ± 0.134 | | -0.008 (-0.096, 0.080) | | 0.023, p = 0.775 | | 0.944 [0.835, 0.984] | | | 0.943 | | 0.031 | | 0.086 |
| Step length α | 100PWS | 0.623 ± 0.131 | 0.606 ± 0.129 | | -0.017 (-0.154, 0.120) | | -0.013, p = 0.916 | | 0.853 [0.638, 0.954] | | | 0.854 | | 0.049 | | 0.137 |
|  | 120PWS | 0.624 ± 0.134 | 0.616 ± 0.145 | | -0.008 (-0.129, 0.114) | | 0.087, p = 0.396 | | 0.904 [0.795, 0.958] | | | 0.905 | | 0.043 | | 0.118 |
|  | 140PWS | 0.668 ± 0.120 | 0.657 ± 0.122 | | -0.011 (-0.126, 0.105) | | 0.017, p = 0.881 | | 0.884 [0.771, 0.936] | | | 0.883 | | 0.041 | | 0.113 |
|  | Self-paced | 0.978 ± 0.157 | 0.952 ± 0.157 | | -0.025 (-0.119, 0.069) | | -0.003, p = 0.963 | | 0.943 [0.878, 0.977] | | | 0.953 | | 0.037 | | 0.103 |
| Step width α | 100PWS | 0.855 ± 0.099 | 0.794 ± 0.124 | | -0.061 (-0.233, 0.112) | | 0.271, p = 0.149 | | 0.612 [0.229, 0.805] | | | 0.712 | | 0.072 | | 0.199 |
|  | 120PWS | 0.791 ± 0.094 | 0.718 ± 0.148 | | -0.074 (-0.284, 0.137) | | 0.521, **p = 0.009** | | 0.539 [0.221, 0.748] | | | 0.690 | | 0.087 | | 0.241 |
|  | 140PWS | 0.766 ± 0.086 | 0.703 ± 0.121 | | -0.063 (-0.264, 0.138) | | 0.421, p = 0.084 | | 0.452 [0.165, 0.715] | | | 0.552 | | 0.080 | | 0.223 |
|  | Self-paced | 0.837 ± 0.118 | 0.777 ± 0.141 | | -0.061 (-0.249, 0.128) | | 0.210, p = 0.247 | | 0.663 [0.348, 0.846] | | | 0.739 | | 0.077 | | 0.212 |
| Body position α | 100PWS | 1.167 ± 0.161 | 1.092 ± 0.151 | | -0.075 (-0.172, 0.022) | | -0.066, p = 0.353 | | 0.854 [0.701, 0.928] | | | 0.952 | | 0.061 | | 0.169 |
|  | 120PWS | 1.129 ± 0.157 | 1.068 ± 0.155 | | -0.061 (-0.125, 0.003) | | -0.012, p = 0.798 | | 0.909 [0.821, 0.949] | | | 0.978 | | 0.047 | | 0.131 |
|  | 140PWS | 1.094 ± 0.149 | 1.038 ± 0.140 | | -0.057 (-0.111, -0.002) | | -0.058, p = 0.174 | | 0.912 [0.840, 0.945] | | | 0.983 | | 0.043 | | 0.119 |
|  | Self-paced | 1.153 ± 0.185 | 1.066 ± 0.179 | | -0.087 (-0.178, 0.003) | | -0.035, p = 0.553 | | 0.869 [0.723, 0.933] | | | 0.968 | | 0.067 | | 0.185 |
| ICC (A, 1) | < 0.50, poor agreement | | | 0.50 – 0.75, moderate agreement | | 0.75 – 0.90, good agreement | | | | | > 0.90 excellent agreement | | | | | |

**Notes:** Values are group means ± SD. Bias and 95% limits of agreement (LoA) were derived from Bland-Altman analysis. Proportional bias p-values reflect regression of inter-system difference on the measurement mean; bold indicates p < 0.05. ICC (A, 1): intraclass correlation coefficient, two-way mixed-effects model, absolute agreement, single measure. SEM: standard error of measurement. MDC: minimal detectable change. PWS: preferred walking speed.

**Supplementary Table S5.** Agreement between marker-based and markerless systems for nonlinear analyses applied on gait joint angles across walking conditions, separately for the generalized-coordinate and Pose2Sim methods.

| **Measure** | **Speed** | | **Vicon**  **(Mean ± SD)** | **Generalized-coordinate method** | | | | | | | | | **Pose2Sim method** | | | | | | | |
| --- | --- | --- | --- | --- | --- | --- | --- | --- | --- | --- | --- | --- | --- | --- | --- | --- | --- | --- | --- | --- |
|  |  |  |  | **Markerless (Mean ± SD)** | **Bias (95% LoA)** | | **Prop. bias**  **(slope, p)** | **ICC (A, 1)** | **r** | **SEM** | | **MDC** | **Markerless (Mean ± SD)** | **Bias (95% LoA)** | **Prop. bias**  **(slope, p)** | | **ICC (A, 1)** | **r** | **SEM** | **MDC** |
| Hip λshort | 100PWS | | 0.287 ± 0.023 | 0.248 ± 0.022 | -0.038 (-0.076, -0.001) | | 0.000, p = 0.999 | 0.272 [0.126, 0.393] | 0,648 | 0,026 | | 0,071 | 0.265 ± 0.024 | -0.022 (-0.056, 0.012) | 0.070, p = 0.699 | | 0.519 [0.341, 0.653] | 0,738 | 0,018 | 0,050 |
|  | 120PWS | | 0.299 ± 0.025 | 0.261 ± 0.022 | -0.038 (-0.073, -0.003) | | -0.169, p = 0.358 | 0.308 [0.096, 0.498] | 0,719 | 0,025 | | 0,069 | 0.276 ± 0.024 | -0.023 (-0.057, 0.012) | -0.028, p = 0.869 | | 0.535 [0.224, 0.736] | 0,752 | 0,018 | 0,051 |
|  | 140PWS | | 0.313 ± 0.025 | 0.270 ± 0.018 | -0.043 (-0.085, -0.001) | | -0.420, p = 0.086 | 0.181 [0.048, 0.277] | 0,552 | 0,028 | | 0,077 | 0.281 ± 0.019 | -0.031 (-0.067, 0.004) | -0.344, p = 0.072 | | 0.342 [0.155, 0.470] | 0,706 | 0,022 | 0,061 |
|  | Self-paced | | 0.316 ± 0.036 | 0.274 ± 0.036 | -0.041 (-0.072, -0.011) | | 0.005, p = 0.958 | 0.545 [0.291, 0.707] | 0,904 | 0,028 | | 0,077 | 0.291 ± 0.040 | -0.024 (-0.059, 0.011) | 0.122, p = 0.265 | | 0.746 [0.468, 0.864] | 0,898 | 0,020 | 0,055 |
| Knee λshort | 100PWS | | 0.652 ± 0.047 | 0.620 ± 0.050 | -0.033 (-0.143, 0.076) | | 0.115, p = 0.717 | 0.301 [0.064, 0.502] | 0,362 | 0,043 | | 0,119 | 0.665 ± 0.053 | 0.010 (-0.101, 0.120) | 0.133, p = 0.677 | | 0.358 [-0.051, 0.633] | 0,355 | 0,039 | 0,109 |
|  | 120PWS | | 0.612 ± 0.044 | 0.583 ± 0.040 | -0.029 (-0.111, 0.053) | | -0.119, p = 0.647 | 0.414 [0.188, 0.647] | 0,506 | 0,034 | | 0,093 | 0.615 ± 0.038 | 0.003 (-0.073, 0.080) | -0.190, p = 0.439 | | 0.552 [0.323, 0.721] | 0,548 | 0,027 | 0,075 |
|  | 140PWS | | 0.609 ± 0.049 | 0.590 ± 0.037 | -0.019 (-0.097, 0.058) | | -0.333, p = 0.134 | 0.549 [0.272, 0.735] | 0,616 | 0,030 | | 0,082 | 0.616 ± 0.037 | 0.007 (-0.062, 0.076) | -0.344, p = 0.076 | | 0.671 [0.396, 0.832] | 0,699 | 0,025 | 0,068 |
|  | Self-paced | | 0.958 ± 0.076 | 0.932 ± 0.072 | -0.034 (-0.166, 0.098) | | -0.191, p = 0.456 | 0.500 [0.277, 0.714] | 0,552 | 0,051 | | 0,141 | 0.984 ± 0.083 | 0.015 (-0.128, 0.158) | -0.089, p = 0.744 | | 0.508 [0.246, 0.724] | 0,507 | 0,051 | 0,142 |
| Ankle λshort | 100PWS | | 1.128 ± 0.071 | 0.858 ± 0.084 | -0.276 (-0.429, -0.124) | | 0.152, p = 0.587 | 0.061 [-0.004, 0.123] | 0,469 | 0,154 | | 0,426 | 0.928 ± 0.083 | -0.207 (-0.354, -0.060) | 0.124, p = 0.644 | | 0.103 [-0.004, 0.202] | 0,499 | 0,121 | 0,336 |
|  | 120PWS | | 1.144 ± 0.052 | 0.890 ± 0.085 | -0.254 (-0.429, -0.080) | | 0.757, **p = 0.029** | 0.027 [-0.029, 0.074] | 0,228 | 0,144 | | 0,400 | 0.949 ± 0.088 | -0.195 (-0.378, -0.013) | 0.821, **p = 0.022** | | 0.036 [-0.042, 0.109] | 0,189 | 0,120 | 0,332 |
|  | 140PWS | | 1.097 ± 0.079 | 0.900 ± 0.085 | -0.197 (-0.334, -0.060) | | 0.085, p = 0.689 | 0.166 [0.070, 0.257] | 0,641 | 0,117 | | 0,325 | 0.941 ± 0.080 | -0.156 (-0.293, -0.019) | 0.009, p = 0.967 | | 0.211 [0.083, 0.335] | 0,613 | 0,099 | 0,274 |
|  | Self-paced | | 1.258 ± 0.105 | 1.026 ± 0.117 | -0.235 (-0.442, -0.028) | | 0.159, p = 0.529 | 0.177 [0.053, 0.280] | 0,563 | 0,148 | | 0,409 | 1.103 ± 0.113 | -0.156 (-0.342, 0.030) | 0.115, p = 0.610 | | 0.320 [0.154, 0.462] | 0,635 | 0,111 | 0,308 |
| Hip ACI | 100PWS | | 0.090 ± 0.011 | 0.081 ± 0.011 | -0.008 (-0.019, 0.002) | | 0.055, p = 0.628 | 0.695 [0.425, 0.831] | 0,889 | 0,006 | | 0,018 | 0.084 ± 0.013 | -0.005 (-0.015, 0.005) | 0.160, p = 0.101 | | 0.831 [0.625, 0.924] | 0,920 | 0,005 | 0,014 |
|  | 120PWS | | 0.096 ± 0.009 | 0.088 ± 0.009 | -0.008 (-0.017, 0.001) | | 0.038, p = 0.754 | 0.629 [0.313, 0.790] | 0,867 | 0,006 | | 0,017 | 0.088 ± 0.010 | -0.007 (-0.018, 0.003) | 0.061, p = 0.639 | | 0.647 [0.327, 0.808] | 0,847 | 0,006 | 0,016 |
|  | 140PWS | | 0.095 ± 0.016 | 0.085 ± 0.014 | -0.010 (-0.021, 0.000) | | -0.110, p = 0.173 | 0.755 [0.556, 0.859] | 0,941 | 0,008 | | 0,022 | 0.087 ± 0.015 | -0.009 (-0.018, 0.000) | -0.064, p = 0.348 | | 0.823 [0.634, 0.905] | 0,957 | 0,007 | 0,018 |
|  | Self-paced | | 0.122 ± 0.018 | 0.116 ± 0.018 | -0.006 (-0.020, 0.008) | | 0.039, p = 0.669 | 0.882 [0.639, 0.954] | 0,928 | 0,006 | | 0,017 | 0.117 ± 0.019 | -0.005 (-0.017, 0.006) | 0.077, p = 0.291 | | 0.915 [0.735, 0.965] | 0,953 | 0,005 | 0,015 |
| Knee ACI | 100PWS | | 0.080 ± 0.010 | 0.080 ± 0.012 | -0.000 (-0.012, 0.011) | | 0.147, p = 0.266 | 0.852 [0.658, 0.926] | 0,854 | 0,004 | | 0,011 | 0.082 ± 0.011 | 0.002 (-0.008, 0.012) | 0.092, p = 0.442 | | 0.859 [0.695, 0.927] | 0,878 | 0,004 | 0,011 |
|  | 120PWS | | 0.089 ± 0.011 | 0.088 ± 0.010 | -0.001 (-0.009, 0.008) | | -0.089, p = 0.337 | 0.920 [0.809, 0.956] | 0,920 | 0,003 | | 0,008 | 0.090 ± 0.012 | 0.002 (-0.009, 0.012) | 0.057, p = 0.589 | | 0.891 [0.738, 0.948] | 0,898 | 0,004 | 0,010 |
|  | 140PWS | | 0.086 ± 0.014 | 0.085 ± 0.013 | -0.001 (-0.010, 0.008) | | -0.078, p = 0.296 | 0.946 [0.856, 0.978] | 0,949 | 0,003 | | 0,009 | 0.087 ± 0.013 | 0.001 (-0.010, 0.011) | -0.075, p = 0.397 | | 0.927 [0.815, 0.966] | 0,927 | 0,004 | 0,010 |
|  | Self-paced | | 0.098 ± 0.013 | 0.101 ± 0.011 | 0.002 (-0.013, 0.016) | | -0.137, p = 0.350 | 0.824 [0.285, 0.981] | 0,830 | 0,005 | | 0,014 | 0.102 ± 0.011 | 0.002 (-0.011, 0.016) | -0.202, p = 0.144 | | 0.826 [0.390, 0.960] | 0,851 | 0,005 | 0,014 |
| Ankle ACI | 100PWS | | 0.061 ± 0.012 | 0.058 ± 0.011 | -0.003 (-0.018, 0.013) | | -0.133, p = 0.420 | 0.761 [0.497, 0.936] | 0,778 | 0,006 | | 0,016 | 0.058 ± 0.010 | -0.003 (-0.017, 0.012) | -0.167, p = 0.278 | | 0.783 [0.597, 0.889] | 0,806 | 0,005 | 0,015 |
|  | 120PWS | | 0.067 ± 0.010 | 0.065 ± 0.009 | -0.001 (-0.011, 0.009) | | -0.109, p = 0.390 | 0.851 [0.674, 0.928] | 0,857 | 0,004 | | 0,010 | 0.064 ± 0.009 | -0.002 (-0.013, 0.008) | -0.134, p = 0.314 | | 0.818 [0.646, 0.898] | 0,845 | 0,004 | 0,011 |
|  | 140PWS | | 0.064 ± 0.011 | 0.064 ± 0.011 | -0.000 (-0.011, 0.011) | | -0.045, p = 0.701 | 0.879 [0.703, 0.941] | 0,875 | 0,004 | | 0,011 | 0.061 ± 0.011 | -0.002 (-0.013, 0.008) | -0.083, p = 0.456 | | 0.872 [0.684, 0.945] | 0,888 | 0,004 | 0,011 |
|  | Self-paced | | 0.066 ± 0.012 | 0.065 ± 0.012 | -0.001 (-0.016, 0.013) | | -0.032, p = 0.836 | 0.812 [0.651, 0.897] | 0,809 | 0,005 | | 0,014 | 0.063 ± 0.011 | -0.003 (-0.019, 0.012) | -0.119, p = 0.483 | | 0.754 [0.493, 0.881] | 0,777 | 0,006 | 0,016 |
| Hip SampEn | 100PWS | | 0.108 ± 0.012 | 0.125 ± 0.017 | 0.017 (-0.007, 0.041) | | 0.462, **p = 0.020** | 0.399 [0.146, 0.575] | 0,704 | 0,013 | | 0,036 | 0.113 ± 0.013 | 0.005 (-0.013, 0.023) | 0.166, p = 0.360 | | 0.692 [0.393, 0.853] | 0,738 | 0,007 | 0,019 |
|  | 120PWS | | 0.111 ± 0.012 | 0.131 ± 0.019 | 0.020 (-0.008, 0.048) | | 0.516, **p = 0.016** | 0.337 [0.167, 0.446] | 0,649 | 0,015 | | 0,042 | 0.119 ± 0.015 | 0.007 (-0.011, 0.026) | 0.239, p = 0.143 | | 0.671 [0.451, 0.811] | 0,776 | 0,008 | 0,023 |
|  | 140PWS | | 0.119 ± 0.015 | 0.138 ± 0.020 | 0.019 (-0.013, 0.051) | | 0.337, p = 0.130 | 0.382 [0.184, 0.510] | 0,615 | 0,016 | | 0,044 | 0.133 ± 0.015 | 0.014 (-0.009, 0.036) | -0.030, p = 0.873 | | 0.505 [0.302, 0.643] | 0,703 | 0,012 | 0,032 |
|  | Self-paced | | 0.112 ± 0.012 | 0.133 ± 0.018 | 0.021 (-0.002, 0.044) | | 0.448, **p = 0.012** | 0.367 [0.218, 0.471] | 0,764 | 0,015 | | 0,041 | 0.119 ± 0.015 | 0.006 (-0.008, 0.020) | 0.215, p = 0.075 | | 0.787 [0.630, 0.874] | 0,881 | 0,006 | 0,017 |
| Knee SampEn | 100PWS | | 0.168 ± 0.029 | 0.145 ± 0.028 | -0.023 (-0.061, 0.015) | | -0.042, p = 0.800 | 0.594 [0.320, 0.760] | 0,775 | 0,020 | | 0,054 | 0.136 ± 0.029 | -0.031 (-0.068, 0.005) | 0.007, p = 0.966 | | 0.511 [0.242, 0.674] | 0,799 | 0,023 | 0,064 |
|  | 120PWS | | 0.199 ± 0.025 | 0.181 ± 0.026 | -0.018 (-0.051, 0.015) | | 0.034, p = 0.831 | 0.631 [0.443, 0.755] | 0,780 | 0,016 | | 0,045 | 0.176 ± 0.024 | -0.022 (-0.058, 0.013) | -0.072, p = 0.687 | | 0.518 [0.267, 0.711] | 0,730 | 0,019 | 0,051 |
|  | 140PWS | | 0.233 ± 0.029 | 0.221 ± 0.028 | -0.012 (-0.058, 0.034) | | -0.033, p = 0.873 | 0.617 [0.353, 0.734] | 0,663 | 0,018 | | 0,049 | 0.213 ± 0.025 | -0.020 (-0.065, 0.025) | -0.188, p = 0.379 | | 0.499 [0.222, 0.703] | 0,638 | 0,020 | 0,055 |
|  | Self-paced | | 0.181 ± 0.034 | 0.161 ± 0.033 | -0.018 (-0.053, 0.016) | | -0.032, p = 0.806 | 0.761 [0.586, 0.874] | 0,865 | 0,017 | | 0,047 | 0.152 ± 0.034 | -0.027 (-0.056, 0.003) | -0.020, p = 0.851 | | 0.693 [0.448, 0.829] | 0,904 | 0,020 | 0,056 |
| Ankle SampEn | 100PWS | | 0.195 ± 0.031 | 0.301 ± 0.049 | 0.103 (-0.013, 0.219) | | 0.898, p = 0.058 | -0.016 [-0.099, 0.052] | -0,073 | 0,066 | | 0,184 | 0.281 ± 0.072 | 0.083 (-0.076, 0.241) | 1.452, **p = < 0.001** | | -0.022 [-0.146, 0.070] | -0,063 | 0,070 | 0,194 |
|  | 120PWS | | 0.194 ± 0.041 | 0.283 ± 0.043 | 0.089 (-0.012, 0.190) | | 0.063, p = 0.857 | 0.079 [-0.036, 0.232] | 0,253 | 0,059 | | 0,163 | 0.249 ± 0.065 | 0.055 (-0.083, 0.192) | 0.725, **p = 0.046** | | 0.116 [-0.107, 0.357] | 0,188 | 0,057 | 0,158 |
|  | 140PWS | | 0.202 ± 0.047 | 0.282 ± 0.039 | 0.080 (-0.023, 0.183) | | -0.282, p = 0.418 | 0.092 [-0.036, 0.256] | 0,252 | 0,056 | | 0,155 | 0.253 ± 0.066 | 0.051 (-0.092, 0.193) | 0.570, p = 0.115 | | 0.141 [-0.056, 0.368] | 0,203 | 0,058 | 0,160 |
|  | Self-paced | | 0.201 ± 0.030 | 0.286 ± 0.041 | 0.084 (-0.019, 0.186) | | 0.697, p = 0.155 | -0.015 [-0.162, 0.097] | -0,057 | 0,056 | | 0,155 | 0.267 ± 0.064 | 0.067 (-0.077, 0.212) | 1.402, **p = 0.001** | | -0.032 [-0.216, 0.106] | -0,077 | 0,062 | 0,171 |
| ICC (A, 1) | | < 0.50, poor agreement | | | | 0.50 – 0.75, moderate agreement | | | | | 0.75 – 0.90, good agreement | | | | | > 0.90 excellent agreement | | | | |

**Notes:** Values are group means ± SD. Bias and 95% limits of agreement (LoA) were derived from Bland-Altman analysis. Proportional bias p-values reflect regression of inter-system difference on the measurement mean; bold indicates p < 0.05. ICC (A, 1): intraclass correlation coefficient, two-way mixed-effects model, absolute agreement, single measure. SEM: standard error of measurement. MDC: minimal detectable change. PWS: preferred walking speed. λshort: maximum Lyapunov exponent. ACI: attractor complexity index. SampEn: sample entropy.

**Supplementary Table S6.** Agreement between marker-based and markerless systems for nonlinear analyses of trunk accelerations across walking conditions.

| **Measure** | **Speed** | **Vicon**  **(Mean ± SD)** | **Markerless (Mean ± SD)** | | **Bias (95% LoA)** | | **Prop. bias**  **(slope, p)** | **ICC (A, 1)** | | **r** | **SEM** | **MDC** |
| --- | --- | --- | --- | --- | --- | --- | --- | --- | --- | --- | --- | --- |
| AP trunk acceleration λshort | 100PWS | 1.188 ± 0.130 | 1.199 ± 0.118 | | 0.011 (-0.186, 0.207) | | -0.117, p = 0.559 | 0.680 [0.432, 0.785] | | 0.676 | 0.069 | 0.192 |
|  | 120PWS | 1.168 ± 0.112 | 1.195 ± 0.112 | | 0.027 (-0.168, 0.222) | | 0.007, p = 0.974 | 0.599 [0.360, 0.728] | | 0.606 | 0.071 | 0.196 |
|  | 140PWS | 1.184 ± 0.105 | 1.267 ± 0.097 | | 0.084 (-0.093, 0.261) | | -0.095, p = 0.673 | 0.452 [0.200, 0.673] | | 0.602 | 0.080 | 0.222 |
|  | Self-paced | 1.313 ± 0.136 | 1.324 ± 0.145 | | 0.010 (-0.185, 0.205) | | 0.072, p = 0.681 | 0.755 [0.437, 0.878] | | 0.750 | 0.069 | 0.190 |
| ML trunk acceleration λshort | 100PWS | 0.874 ± 0.113 | 1.220 ± 0.115 | | 0.345 (0.135, 0.556) | | 0.027, p = 0.910 | 0.100 [0.013, 0.190] | | 0.558 | 0.197 | 0.547 |
|  | 120PWS | 0.952 ± 0.110 | 1.349 ± 0.100 | | 0.397 (0.197, 0.596) | | -0.122, p = 0.625 | 0.065 [0.003, 0.128] | | 0.533 | 0.218 | 0.605 |
|  | 140PWS | 0.952 ± 0.121 | 1.420 ± 0.103 | | 0.467 (0.269, 0.665) | | -0.207, p = 0.360 | 0.062 [0.011, 0.108] | | 0.604 | 0.253 | 0.701 |
|  | Self-paced | 1.037 ± 0.158 | 1.371 ± 0.149 | | 0.334 (0.108, 0.560) | | -0.072, p = 0.700 | 0.214 [0.085, 0.309] | | 0.720 | 0.201 | 0.558 |
| VT trunk acceleration λshort | 100PWS | 1.118 ± 0.111 | 1.306 ± 0.127 | | 0.187 (0.030, 0.345) | | 0.158, p = 0.323 | 0.348 [0.203, 0.467] | | 0.781 | 0.122 | 0.338 |
|  | 120PWS | 1.183 ± 0.117 | 1.349 ± 0.124 | | 0.166 (0.021, 0.312) | | 0.068, p = 0.639 | 0.418 [0.271, 0.542] | | 0.813 | 0.111 | 0.309 |
|  | 140PWS | 1.191 ± 0.097 | 1.372 ± 0.135 | | 0.181 (-0.009, 0.372) | | 0.382, p = 0.052 | 0.303 [0.118, 0.462] | | 0.694 | 0.124 | 0.343 |
|  | Self-paced | 1.202 ± 0.128 | 1.391 ± 0.170 | | 0.189 (-0.003, 0.381) | | 0.305, p = 0.045 | 0.443 [0.223, 0.630] | | 0.818 | 0.132 | 0.365 |
| AP trunk acceleration ACI | 100PWS | 0.073 ± 0.011 | 0.067 ± 0.011 | | -0.007 (-0.017, 0.004) | | -0.026, p = 0.812 | 0.767 [0.488, 0.873] | | 0.891 | 0.006 | 0.016 |
|  | 120PWS | 0.080 ± 0.008 | 0.075 ± 0.007 | | -0.005 (-0.014, 0.004) | | -0.158, p = 0.272 | 0.680 [0.386, 0.827] | | 0.820 | 0.005 | 0.013 |
|  | 140PWS | 0.078 ± 0.013 | 0.074 ± 0.013 | | -0.004 (-0.015, 0.007) | | -0.007, p = 0.944 | 0.868 [0.718, 0.943] | | 0.906 | 0.005 | 0.013 |
|  | Self-paced | 0.072 ± 0.016 | 0.069 ± 0.016 | | -0.003 (-0.015, 0.008) | | -0.022, p = 0.804 | 0.912 [0.791, 0.966] | | 0.929 | 0.005 | 0.013 |
| ML trunk acceleration ACI | 100PWS | 0.054 ± 0.011 | 0.051 ± 0.012 | | -0.002 (-0.018, 0.013) | | 0.068, p = 0.695 | 0.734 [0.441, 0.887] | | 0.744 | 0.006 | 0.016 |
|  | 120PWS | 0.060 ± 0.011 | 0.058 ± 0.010 | | -0.002 (-0.016, 0.012) | | -0.048, p = 0.771 | 0.766 [0.555, 0.883] | | 0.770 | 0.005 | 0.014 |
|  | 140PWS | 0.056 ± 0.011 | 0.056 ± 0.011 | | -0.000 (-0.015, 0.014) | | 0.023, p = 0.885 | 0.782 [0.538, 0.886] | | 0.775 | 0.005 | 0.014 |
|  | Self-paced | 0.061 ± 0.009 | 0.066 ± 0.012 | | 0.004 (-0.012, 0.021) | | 0.402, p = 0.038 | 0.627 [0.406, 0.787] | | 0.717 | 0.006 | 0.018 |
| VT trunk acceleration ACI | 100PWS | 0.069 ± 0.013 | 0.063 ± 0.013 | | -0.006 (-0.015, 0.003) | | -0.016, p = 0.837 | 0.844 [0.597, 0.906] | | 0.940 | 0.005 | 0.015 |
|  | 120PWS | 0.081 ± 0.009 | 0.077 ± 0.009 | | -0.004 (-0.011, 0.004) | | -0.009, p = 0.923 | 0.842 [0.647, 0.913] | | 0.910 | 0.004 | 0.010 |
|  | 140PWS | 0.081 ± 0.012 | 0.077 ± 0.012 | | -0.003 (-0.011, 0.004) | | 0.026, p = 0.734 | 0.910 [0.734, 0.966] | | 0.945 | 0.004 | 0.010 |
|  | Self-paced | 0.078 ± 0.018 | 0.066 ± 0.021 | | -0.012 (-0.033, 0.009) | | 0.143, p = 0.272 | 0.719 [0.311, 0.873] | | 0.857 | 0.011 | 0.030 |
| AP trunk acceleration SampEn | 100PWS | 0.190 ± 0.018 | 0.295 ± 0.027 | | 0.105 (0.036, 0.175) | | 0.975, p = 0.074 | -0.019 [-0.054, 0.011] | | -0.240 | 0.058 | 0.161 |
|  | 120PWS | 0.177 ± 0.020 | 0.292 ± 0.030 | | 0.115 (0.043, 0.188) | | 0.781, p = 0.083 | -0.003 [-0.043, 0.026] | | -0.035 | 0.064 | 0.177 |
|  | 140PWS | 0.166 ± 0.019 | 0.284 ± 0.024 | | 0.118 (0.061, 0.174) | | 0.427, p = 0.283 | 0.007 [-0.014, 0.026] | | 0.117 | 0.063 | 0.175 |
|  | Self-paced | 0.195 ± 0.022 | 0.298 ± 0.026 | | 0.103 (0.033, 0.174) | | 0.327, p = 0.523 | -0.010 [-0.058, 0.030] | | -0.101 | 0.058 | 0.160 |
| ML trunk acceleration SampEn | 100PWS | 0.307 ± 0.054 | 0.208 ± 0.039 | | -0.099 (-0.208, 0.010) | | -0.488, p = 0.132 | 0.093 [-0.012, 0.218] | | 0.313 | 0.065 | 0.180 |
|  | 120PWS | 0.338 ± 0.053 | 0.263 ± 0.048 | | -0.075 (-0.196, 0.046) | | -0.155, p = 0.652 | 0.130 [-0.069, 0.356] | | 0.267 | 0.059 | 0.163 |
|  | 140PWS | 0.365 ± 0.056 | 0.298 ± 0.056 | | -0.066 (-0.206, 0.074) | | -0.018, p = 0.963 | 0.108 [-0.199, 0.447] | | 0.180 | 0.061 | 0.169 |
|  | Self-paced | 0.315 ± 0.059 | 0.225 ± 0.064 | | -0.089 (-0.218, 0.039) | | 0.105, p = 0.720 | 0.211 [0.009, 0.431] | | 0.430 | 0.067 | 0.186 |
| VT trunk acceleration SampEn | 100PWS | 0.318 ± 0.032 | 0.307 ± 0.042 | | -0.011 (-0.098, 0.076) | | 0.435, p = 0.180 | 0.297 [-0.243, 0.619] | | 0.313 | 0.031 | 0.087 |
|  | 120PWS | 0.287 ± 0.047 | 0.279 ± 0.036 | | -0.008 (-0.100, 0.084) | | -0.365, p = 0.220 | 0.384 [-0.108, 0.664] | | 0.392 | 0.033 | 0.091 |
|  | 140PWS | 0.254 ± 0.039 | 0.279 ± 0.037 | | 0.025 (-0.039, 0.089) | | -0.052, p = 0.808 | 0.527 [0.306, 0.685] | | 0.632 | 0.027 | 0.075 |
|  | Self-paced | 0.339 ± 0.034 | 0.323 ± 0.047 | | -0.016 (-0.124, 0.091) | | 0.544, p = 0.185 | 0.098 [-0.356, 0.557] | | 0.107 | 0.039 | 0.109 |
| ICC (A, 1) | < 0.50, poor agreement | | | 0.50 – 0.75, moderate agreement | | 0.75 – 0.90, good agreement | | | > 0.90 excellent agreement | | | |

**Notes:** Values are group means ± SD. Bias and 95% limits of agreement (LoA) were derived from Bland-Altman analysis. Proportional bias p-values reflect regression of inter-system difference on the measurement mean; bold indicates p < 0.05. ICC (A, 1): intraclass correlation coefficient, two-way mixed-effects model, absolute agreement, single measure. SEM: standard error of measurement. MDC: minimal detectable change. PWS: preferred walking speed. λshort: maximum Lyapunov exponent. ACI: attractor complexity index. SampEn: sample entropy. AP: anteroposterior. ML: mediolateral. VT: vertical.

**Supplementary Table S7.** Agreement between marker-based and markerless systems for nonlinear analyses of trunk positions across walking conditions.

| **Measure** | **Speed** | **Vicon**  **(Mean ± SD)** | **Markerless (Mean ± SD)** | | **Bias (95% LoA)** | | **Prop. bias**  **(slope, p)** | **ICC (A, 1)** | | **r** | **SEM** | **MDC** |
| --- | --- | --- | --- | --- | --- | --- | --- | --- | --- | --- | --- | --- |
| AP trunk position λshort | 100PWS | 1.737 ± 0.104 | 1.726 ± 0.094 | | -0.011 (-0.095, 0.073) | | -0.101, p = 0.294 | 0.909 [0.773, 0.965] | | 0.915 | 0.030 | 0.084 |
|  | 120PWS | 1.785 ± 0.087 | 1.774 ± 0.097 | | -0.011 (-0.099, 0.076) | | 0.124, p = 0.253 | 0.885 [0.610, 0.949] | | 0.893 | 0.032 | 0.088 |
|  | 140PWS | 1.639 ± 0.134 | 1.670 ± 0.117 | | 0.031 (-0.133, 0.194) | | -0.154, p = 0.316 | 0.774 [0.490, 0.899] | | 0.797 | 0.061 | 0.168 |
|  | Self-paced | 1.881 ± 0.105 | 1.862 ± 0.116 | | -0.020 (-0.097, 0.057) | | 0.105, p = 0.189 | 0.928 [0.851, 0.961] | | 0.945 | 0.030 | 0.084 |
| ML trunk position λshort | 100PWS | 1.336 ± 0.159 | 1.347 ± 0.167 | | 0.011 (-0.298, 0.320) | | 0.058, p = 0.812 | 0.565 [0.295, 0.787] | | 0.556 | 0.109 | 0.302 |
|  | 120PWS | 1.276 ± 0.169 | 1.276 ± 0.180 | | -0.000 (-0.197, 0.197) | | 0.068, p = 0.609 | 0.849 [0.691, 0.930] | | 0.844 | 0.069 | 0.191 |
|  | 140PWS | 1.286 ± 0.158 | 1.239 ± 0.154 | | -0.047 (-0.323, 0.228) | | -0.032, p = 0.885 | 0.595 [0.270, 0.853] | | 0.612 | 0.101 | 0.281 |
|  | Self-paced | 1.915 ± 0.135 | 1.832 ± 0.115 | | -0.083 (-0.181, 0.014) | | -0.162, p = 0.065 | 0.767 [0.585, 0.862] | | 0.937 | 0.065 | 0.179 |
| VT trunk position λshort | 100PWS | 2.004 ± 0.122 | 1.749 ± 0.120 | | -0.255 (-0.397, -0.113) | | -0.015, p = 0.915 | 0.266 [0.120, 0.391] | | 0.829 | 0.152 | 0.422 |
|  | 120PWS | 2.675 ± 0.133 | 2.388 ± 0.144 | | -0.287 (-0.561, -0.012) | | 0.101, p = 0.694 | 0.170 [0.017, 0.317] | | 0.514 | 0.184 | 0.509 |
|  | 140PWS | 2.741 ± 0.132 | 2.524 ± 0.137 | | -0.217 (-0.499, 0.064) | | 0.045, p = 0.870 | 0.205 [-0.009, 0.396] | | 0.456 | 0.156 | 0.432 |
|  | Self-paced | 2.184 ± 0.151 | 1.964 ± 0.165 | | -0.219 (-0.385, -0.054) | | 0.093, p = 0.455 | 0.454 [0.212, 0.609] | | 0.868 | 0.144 | 0.399 |
| AP trunk position ACI | 100PWS | 0.015 ± 0.013 | 0.016 ± 0.013 | | 0.001 (-0.006, 0.008) | | -0.011, p = 0.864 | 0.957 [0.842, 0.985] | | 0.960 | 0.003 | 0.007 |
|  | 120PWS | 0.014 ± 0.009 | 0.013 ± 0.008 | | -0.001 (-0.009, 0.008) | | -0.147, p = 0.215 | 0.870 [0.713, 0.940] | | 0.875 | 0.003 | 0.008 |
|  | 140PWS | 0.008 ± 0.011 | 0.010 ± 0.012 | | 0.002 (-0.005, 0.008) | | 0.023, p = 0.715 | 0.950 [0.877, 0.981] | | 0.961 | 0.003 | 0.007 |
|  | Self-paced | 0.025 ± 0.013 | 0.026 ± 0.013 | | 0.001 (-0.005, 0.008) | | 0.027, p = 0.660 | 0.963 [0.897, 0.983] | | 0.967 | 0.002 | 0.007 |
| ML trunk position ACI | 100PWS | 0.049 ± 0.017 | 0.049 ± 0.017 | | -0.000 (-0.007, 0.006) | | 0.006, p = 0.882 | 0.984 [0.953, 0.992] | | 0.983 | 0.002 | 0.006 |
|  | 120PWS | 0.059 ± 0.019 | 0.059 ± 0.019 | | 0.000 (-0.006, 0.007) | | 0.012, p = 0.762 | 0.985 [0.966, 0.994] | | 0.985 | 0.002 | 0.007 |
|  | 140PWS | 0.063 ± 0.019 | 0.064 ± 0.019 | | 0.000 (-0.007, 0.007) | | 0.012, p = 0.769 | 0.984 [0.963, 0.994] | | 0.983 | 0.002 | 0.007 |
|  | Self-paced | 0.016 ± 0.017 | 0.015 ± 0.016 | | -0.002 (-0.012, 0.009) | | -0.026, p = 0.728 | 0.949 [0.826, 0.988] | | 0.951 | 0.004 | 0.011 |
| VT trunk position ACI | 100PWS | 0.067 ± 0.012 | 0.058 ± 0.012 | | -0.009 (-0.017, -0.000) | | 0.019, p = 0.796 | 0.771 [0.474, 0.868] | | 0.947 | 0.006 | 0.017 |
|  | 120PWS | 0.075 ± 0.009 | 0.071 ± 0.009 | | -0.005 (-0.014, 0.004) | | 0.097, p = 0.407 | 0.772 [0.560, 0.885] | | 0.876 | 0.004 | 0.012 |
|  | 140PWS | 0.074 ± 0.013 | 0.069 ± 0.011 | | -0.005 (-0.016, 0.005) | | -0.142, p = 0.149 | 0.825 [0.590, 0.917] | | 0.913 | 0.005 | 0.014 |
|  | Self-paced | 0.079 ± 0.016 | 0.066 ± 0.017 | | -0.012 (-0.033, 0.008) | | 0.123, p = 0.416 | 0.639 [0.265, 0.839] | | 0.812 | 0.011 | 0.030 |
| AP trunk position SampEn | 100PWS | 0.076 ± 0.026 | 0.080 ± 0.026 | | 0.004 (-0.002, 0.010) | | 0.001, p = 0.980 | 0.984 [0.958, 0.993] | | 0.994 | 0.003 | 0.009 |
|  | 120PWS | 0.079 ± 0.030 | 0.085 ± 0.031 | | 0.006 (-0.001, 0.013) | | 0.008, p = 0.752 | 0.977 [0.949, 0.988] | | 0.994 | 0.005 | 0.013 |
|  | 140PWS | 0.076 ± 0.031 | 0.085 ± 0.033 | | 0.009 (-0.001, 0.020) | | 0.065, p = 0.062 | 0.948 [0.894, 0.972] | | 0.989 | 0.007 | 0.020 |
|  | Self-paced | 0.083 ± 0.028 | 0.088 ± 0.029 | | 0.005 (-0.001, 0.011) | | 0.021, p = 0.377 | 0.982 [0.967, 0.990] | | 0.995 | 0.004 | 0.011 |
| ML trunk position SampEn | 100PWS | 0.037 ± 0.010 | 0.041 ± 0.011 | | 0.004 (-0.002, 0.010) | | 0.035, p = 0.603 | 0.899 [0.729, 0.953] | | 0.957 | 0.003 | 0.010 |
|  | 120PWS | 0.032 ± 0.012 | 0.037 ± 0.014 | | 0.005 (-0.003, 0.013) | | 0.115, p = 0.082 | 0.903 [0.755, 0.964] | | 0.961 | 0.004 | 0.012 |
|  | 140PWS | 0.036 ± 0.011 | 0.042 ± 0.012 | | 0.005 (-0.002, 0.012) | | 0.087, p = 0.186 | 0.873 [0.738, 0.933] | | 0.960 | 0.004 | 0.012 |
|  | Self-paced | 0.058 ± 0.016 | 0.065 ± 0.018 | | 0.007 (-0.005, 0.020) | | 0.120, p = 0.160 | 0.854 [0.716, 0.907] | | 0.938 | 0.007 | 0.018 |
| VT trunk position SampEn | 100PWS | 0.293 ± 0.020 | 0.304 ± 0.022 | | 0.011 (-0.023, 0.046) | | 0.137, p = 0.496 | 0.593 [0.194, 0.834] | | 0.670 | 0.014 | 0.039 |
|  | 120PWS | 0.288 ± 0.021 | 0.306 ± 0.024 | | 0.018 (-0.003, 0.039) | | 0.135, p = 0.197 | 0.696 [0.444, 0.837] | | 0.902 | 0.014 | 0.038 |
|  | 140PWS | 0.286 ± 0.022 | 0.309 ± 0.028 | | 0.023 (-0.002, 0.048) | | 0.246, **p = 0.026** | 0.621 [0.384, 0.754] | | 0.898 | 0.017 | 0.047 |
|  | Self-paced | 0.303 ± 0.021 | 0.320 ± 0.024 | | 0.017 (-0.025, 0.058) | | 0.137, p = 0.563 | 0.471 [0.165, 0.732] | | 0.589 | 0.018 | 0.049 |
| ICC (A, 1) | < 0.50, poor agreement | | | 0.50 – 0.75, moderate agreement | | 0.75 – 0.90, good agreement | | | > 0.90 excellent agreement | | | |

**Notes:** Values are group means ± SD. Bias and 95% limits of agreement (LoA) were derived from Bland-Altman analysis. Proportional bias p-values reflect regression of inter-system difference on the measurement mean; bold indicates p < 0.05. ICC (A, 1): intraclass correlation coefficient, two-way mixed-effects model, absolute agreement, single measure. SEM: standard error of measurement. MDC: minimal detectable change. PWS: preferred walking speed. λshort: maximum Lyapunov exponent. ACI: attractor complexity index. SampEn: sample entropy. AP: anteroposterior. ML: mediolateral. VT: vertical.

**Supplementary Table S8**. Agreement between marker-based and markerless systems for nonlinear analyses of trunk velocities across walking conditions.

| **Measure** | **Speed** | **Vicon**  **(Mean ± SD)** | **Markerless (Mean ± SD)** | | **Bias (95% LoA)** | | **Prop. bias**  **(slope, p)** | **ICC (A, 1)** | | **r** | **SEM** | **MDC** |
| --- | --- | --- | --- | --- | --- | --- | --- | --- | --- | --- | --- | --- |
| AP trunk velocity λshort | 100PWS | 1.464 ± 0.101 | 1.534 ± 0.124 | | 0.070 (-0.154, 0.294) | | 0.263, p = 0.299 | 0.444 [0.112, 0.700] | | 0.524 | 0.089 | 0.248 |
|  | 120PWS | 1.883 ± 0.091 | 2.045 ± 0.198 | | 0.162 (-0.245, 0.569) | | 1.146, **p = 0.001** | 0.090 [-0.110, 0.370] | | 0.176 | 0.168 | 0.466 |
|  | 140PWS | 1.767 ± 0.110 | 2.017 ± 0.149 | | 0.250 (-0.129, 0.630) | | 0.619, p = 0.182 | -0.017 [-0.137, 0.155] | | -0.049 | 0.185 | 0.512 |
|  | Self-paced | 1.172 ± 0.082 | 1.246 ± 0.088 | | 0.074 (-0.097, 0.244) | | 0.093, p = 0.725 | 0.380 [0.096, 0.630] | | 0.507 | 0.074 | 0.205 |
| ML trunk velocity λshort | 100PWS | 2.209 ± 0.192 | 1.757 ± 0.145 | | -0.452 (-0.820, -0.085) | | -0.387, p = 0.170 | 0.097 [0.017, 0.175] | | 0.439 | 0.272 | 0.754 |
|  | 120PWS | 2.153 ± 0.205 | 1.782 ± 0.191 | | -0.371 (-0.761, 0.020) | | -0.093, p = 0.716 | 0.195 [0.037, 0.370] | | 0.518 | 0.246 | 0.682 |
|  | 140PWS | 2.121 ± 0.175 | 1.894 ± 0.155 | | -0.227 (-0.557, 0.103) | | -0.157, p = 0.544 | 0.268 [0.086, 0.459] | | 0.507 | 0.173 | 0.480 |
|  | Self-paced | 2.252 ± 0.189 | 1.910 ± 0.146 | | -0.341 (-0.684, 0.001) | | -0.335, p = 0.209 | 0.167 [0.015, 0.313] | | 0.506 | 0.222 | 0.614 |
| VT trunk velocity λshort | 100PWS | 2.134 ± 0.181 | 2.078 ± 0.208 | | -0.056 (-0.313, 0.201) | | 0.156, p = 0.315 | 0.763 [0.598, 0.848] | | 0.793 | 0.097 | 0.269 |
|  | 120PWS | 2.051 ± 0.175 | 1.961 ± 0.198 | | -0.089 (-0.362, 0.184) | | 0.142, p = 0.417 | 0.670 [0.427, 0.823] | | 0.741 | 0.112 | 0.310 |
|  | 140PWS | 1.965 ± 0.132 | 1.911 ± 0.172 | | -0.054 (-0.354, 0.246) | | 0.336, p = 0.174 | 0.506 [0.189, 0.713] | | 0.542 | 0.111 | 0.307 |
|  | Self-paced | 2.191 ± 0.202 | 2.200 ± 0.260 | | 0.009 (-0.266, 0.284) | | 0.270, **p = 0.046** | 0.834 [0.639, 0.927] | | 0.854 | 0.096 | 0.267 |
| AP trunk velocity ACI | 100PWS | 0.040 ± 0.012 | 0.041 ± 0.012 | | 0.000 (-0.006, 0.007) | | 0.054, p = 0.386 | 0.964 [0.905, 0.987] | | 0.964 | 0.002 | 0.006 |
|  | 120PWS | 0.044 ± 0.014 | 0.044 ± 0.013 | | -0.000 (-0.008, 0.008) | | -0.046, p = 0.485 | 0.959 [0.907, 0.980] | | 0.959 | 0.003 | 0.008 |
|  | 140PWS | 0.038 ± 0.010 | 0.040 ± 0.011 | | 0.002 (-0.006, 0.009) | | 0.025, p = 0.759 | 0.931 [0.859, 0.963] | | 0.939 | 0.003 | 0.008 |
|  | Self-paced | 0.064 ± 0.014 | 0.065 ± 0.014 | | 0.001 (-0.005, 0.007) | | 0.003, p = 0.959 | 0.975 [0.946, 0.987] | | 0.977 | 0.002 | 0.006 |
| ML trunk velocity ACI | 100PWS | 0.060 ± 0.011 | 0.057 ± 0.011 | | -0.003 (-0.013, 0.006) | | -0.034, p = 0.725 | 0.883 [0.729, 0.939] | | 0.916 | 0.004 | 0.011 |
|  | 120PWS | 0.068 ± 0.008 | 0.066 ± 0.008 | | -0.002 (-0.008, 0.004) | | -0.009, p = 0.917 | 0.920 [0.794, 0.963] | | 0.936 | 0.002 | 0.007 |
|  | 140PWS | 0.063 ± 0.013 | 0.062 ± 0.012 | | -0.001 (-0.009, 0.007) | | -0.016, p = 0.824 | 0.946 [0.885, 0.975] | | 0.950 | 0.003 | 0.008 |
|  | Self-paced | 0.059 ± 0.015 | 0.058 ± 0.015 | | -0.002 (-0.012, 0.009) | | 0.013, p = 0.869 | 0.940 [0.833, 0.982] | | 0.942 | 0.004 | 0.010 |
| VT trunk velocity ACI | 100PWS | 0.076 ± 0.013 | 0.065 ± 0.014 | | -0.011 (-0.021, 0.000) | | 0.071, p = 0.447 | 0.703 [0.376, 0.814] | | 0.920 | 0.008 | 0.022 |
|  | 120PWS | 0.086 ± 0.009 | 0.078 ± 0.009 | | -0.008 (-0.014, -0.001) | | -0.024, p = 0.782 | 0.695 [0.476, 0.792] | | 0.932 | 0.005 | 0.015 |
|  | 140PWS | 0.084 ± 0.012 | 0.077 ± 0.011 | | -0.007 (-0.014, 0.000) | | -0.080, p = 0.240 | 0.820 [0.586, 0.915] | | 0.957 | 0.005 | 0.015 |
|  | Self-paced | 0.086 ± 0.017 | 0.071 ± 0.019 | | -0.015 (-0.038, 0.008) | | 0.165, p = 0.277 | 0.609 [0.211, 0.803] | | 0.811 | 0.012 | 0.034 |
| AP trunk velocity SampEn | 100PWS | 0.192 ± 0.020 | 0.174 ± 0.013 | | -0.018 (-0.050, 0.014) | | -0.547, **p = 0.014** | 0.367 [0.103, 0.623] | | 0.624 | 0.016 | 0.043 |
|  | 120PWS | 0.213 ± 0.023 | 0.199 ± 0.015 | | -0.015 (-0.057, 0.028) | | -0.600, **p = 0.027** | 0.340 [-0.002, 0.727] | | 0.466 | 0.017 | 0.048 |
|  | 140PWS | 0.239 ± 0.028 | 0.223 ± 0.018 | | -0.016 (-0.076, 0.043) | | -0.717, **p = 0.038** | 0.177 [-0.149, 0.697] | | 0.235 | 0.023 | 0.064 |
|  | Self-paced | 0.196 ± 0.019 | 0.182 ± 0.023 | | -0.014 (-0.041, 0.013) | | 0.164, p = 0.287 | 0.664 [0.421, 0.797] | | 0.805 | 0.013 | 0.036 |
| ML trunk velocity SampEn | 100PWS | 0.255 ± 0.021 | 0.302 ± 0.032 | | 0.047 (0.005, 0.089) | | 0.492, **p = 0.004** | 0.289 [0.108, 0.465] | | 0.768 | 0.030 | 0.085 |
|  | 120PWS | 0.240 ± 0.024 | 0.306 ± 0.031 | | 0.066 (0.013, 0.120) | | 0.308, p = 0.200 | 0.150 [0.048, 0.250] | | 0.563 | 0.040 | 0.112 |
|  | 140PWS | 0.232 ± 0.024 | 0.306 ± 0.024 | | 0.073 (0.026, 0.121) | | -0.014, p = 0.955 | 0.094 [0.027, 0.153] | | 0.516 | 0.042 | 0.117 |
|  | Self-paced | 0.265 ± 0.024 | 0.314 ± 0.032 | | 0.049 (0.002, 0.097) | | 0.327, p = 0.114 | 0.267 [0.107, 0.417] | | 0.676 | 0.032 | 0.090 |
| VT trunk velocity SampEn | 100PWS | 0.263 ± 0.036 | 0.312 ± 0.030 | | 0.049 (-0.028, 0.126) | | -0.302, p = 0.344 | 0.162 [-0.050, 0.390] | | 0.332 | 0.038 | 0.105 |
|  | 120PWS | 0.276 ± 0.029 | 0.304 ± 0.024 | | 0.028 (-0.033, 0.089) | | -0.310, p = 0.329 | 0.219 [-0.033, 0.517] | | 0.336 | 0.027 | 0.074 |
|  | 140PWS | 0.285 ± 0.021 | 0.299 ± 0.025 | | 0.014 (-0.029, 0.058) | | 0.250, p = 0.297 | 0.475 [0.062, 0.747] | | 0.564 | 0.018 | 0.049 |
|  | Self-paced | 0.287 ± 0.035 | 0.321 ± 0.027 | | 0.034 (-0.048, 0.116) | | -0.428, p = 0.281 | 0.092 [-0.202, 0.427] | | 0.145 | 0.034 | 0.095 |
| ICC (A, 1) | < 0.50, poor agreement | | | 0.50 – 0.75, moderate agreement | | 0.75 – 0.90, good agreement | | | > 0.90 excellent agreement | | | |

**Notes:** Values are group means ± SD. Bias and 95% limits of agreement (LoA) were derived from Bland-Altman analysis. Proportional bias p-values reflect regression of inter-system difference on the measurement mean; bold indicates p < 0.05. ICC (A, 1): intraclass correlation coefficient, two-way mixed-effects model, absolute agreement, single measure. SEM: standard error of measurement. MDC: minimal detectable change. PWS: preferred walking speed. λshort: maximum Lyapunov exponent. ACI: attractor complexity index. SampEn: sample entropy. AP: anteroposterior. ML: mediolateral. VT: vertical.

**Supplementary Table S9.** Effect of walking speed on inter-system agreement. One-way repeated-measures ANOVAs on inter-system differences (markerless minus Vicon) with walking speed as a four-level within-subject factor.

| **Measure** | **Mauchly W (p)** | **εGG** | **F(df1, df2)** | **p** | **pFDR** | **η²p** | **Bias 100%** | **Bias 120%** | **Bias 140%** | **Bias SP** | **Post-hoc pairs** |
| --- | --- | --- | --- | --- | --- | --- | --- | --- | --- | --- | --- |
| **Spatiotemporal parameters (mean)** | | | | | | | | | | | |
| Step time (s) | 0.220 (0.000) | 0.547 | F(3,60) = 3.21† | 0.062 | 0.127 | 0.138 | 0.0000 | 0.0000 | 0.0000 | 0.0000 | - |
| Cadence (steps/min) | 0.382 (0.003) | 0.656 | F(3,60) = 3.52† | 0.040 | 0.090 | 0.150 | -0.0037 | -0.0030 | -0.0026 | -0.0001 | - |
| Double-support time (s) | 0.472 (0.015) | 0.696 | F(3,60) = 0.32† | 0.740 | 0.812 | 0.016 | 0.0006 | -0.0008 | 0.0000 | 0.0005 | - |
| Single-support time (s) | 0.774 (0.443) | - | F(3,60) = 0.80 | 0.496 | 0.651 | 0.039 | -0.0012 | 0.0003 | -0.0003 | -0.0011 | - |
| Step speed (m/s) | 0.873 (0.771) | - | F(3,60) = 34.30 | < 0.001 | **< 0.001*** | 0.632 | 0.0206 | 0.0281 | 0.0374 | 0.0245 | 100% - 120%  100% - 140%  120% - 140%  120% - SP  140% - SP |
| Step length (m) | 0.847 (0.683) | - | F(3,60) = 19.61 | < 0.001 | **< 0.001*** | 0.495 | 0.0117 | 0.0147 | 0.0183 | 0.0132 | 100% - 120%  100% - 140%  120% - 140%  140% - SP |
| Step width (m) | 0.360 (0.002) | 0.661 | F(3,60) = 4.33† | 0.020 | **0.048*** | 0.178 | 0.0071 | 0.0065 | 0.0056 | 0.0067 | 140% - SP |
| **Spatiotemporal parameters (SD)** | | | | | | | | | | | |
| Step time SD (s) | 0.344 (0.001) | 0.704 | F(3,60) = 1.43† | 0.251 | 0.376 | 0.067 | 0.0002 | 0.0006 | 0.0003 | 0.0002 | - |
| Step speed SD (m/s) | 0.570 (0.061) | - | F(3,60) = 0.70 | 0.556 | 0.673 | 0.034 | 0.0029 | 0.0043 | 0.0037 | 0.0029 | - |
| Step length SD (m) | 0.397 (0.004) | 0.633 | F(3,60) = 0.23† | 0.781 | 0.837 | 0.012 | 0.0015 | 0.0017 | 0.0016 | 0.0015 | - |
| Step width SD (m) | 0.553 (0.050) | 0.802 | F(3,60) = 3.00† | 0.050 | 0.107 | 0.131 | 0.0008 | 0.0021 | 0.0021 | 0.0008 | - |
| Double-support time SD (s) | 0.770 (0.427) | - | F(3,60) = 9.37 | < 0.001 | **< 0.001*** | 0.319 | 0,0004 | 0,0002 | 0,0001 | 0,0009 | 100% - SP  120% - SP  140% - SP |
| Single-support time SD (s) | 0.081 (0.000) | 0.423 | F(3,60) = 3.00† | 0.087 | 0.163 | 0.131 | 0.0007 | 0.0002 | 0.0001 | 0.0000 | - |
| **Spatiotemporal parameters (DFA, α)** | | | | | | | | | | | |
| Step time α | 0.562 (0.056) | - | F(3,60) = 1.52 | 0.219 | 0.340 | 0.071 | -0.0079 | -0.0257 | -0.0151 | -0.0076 | - |
| Step length α | 0.517 (0.030) | 0.681 | F(3,60) = 0.70† | 0.506 | 0.651 | 0.034 | -0.0169 | -0.0078 | -0.0105 | -0.0253 | - |
| Step speed α | 0.347 (0.001) | 0.586 | F(3,60) = 0.18† | 0.811 | 0.849 | 0.009 | -0.0079 | -0.0220 | -0.0056 | -0.0151 | - |
| Step width α | 0.709 (0.265) | - | F(3,60) = 0.06 | 0.982 | 0.982 | 0.003 | -0.0607 | -0.0736 | -0.0627 | -0.0607 | - |
| Body position α | 0.554 (0.051) | - | F(3,60) = 5.53 | 0.002 | **0.010*** | 0.216 | -0.0750 | -0.0610 | -0.0565 | -0.0871 | 140% - SP |
| **Joint angles - Generalized-coordinate method** | | | | | | | | | | | |
| Hip LDS | 0.851 (0.721) | - | F(3,57) = 1.90 | 0.140 | 0.233 | 0.091 | -0.0385 | -0.0380 | -0.0432 | -0.0415 | - |
| Knee LDS | 0.492 (0.037) | 0.664 | F(3,54) = 2.45† | 0.101 | 0.174 | 0.120 | -0.0333 | -0.0290 | -0.0195 | -0.0341 | - |
| Ankle LDS | 0.710 (0.333) | - | F(3,54) = 12.12 | < 0.001 | **< 0.001*** | 0.402 | -0.2765 | -0.2544 | -0.1969 | -0.2348 | 100% - 140%  120% - 140%  140% - SP |
| Hip ACI | 0.666 (0.207) | - | F(3,57) = 4.54 | 0.006 | **0.022*** | 0.193 | -0.0083 | -0.0081 | -0.0105 | -0.0060 | - |
| Knee ACI | 0.689 (0.285) | - | F(3,54) = 0.68 | 0.568 | 0.673 | 0.036 | -0.0002 | -0.0005 | -0.0010 | 0.0015 | - |
| Ankle ACI | 0.608 (0.139) | - | F(3,54) = 1.09 | 0.362 | 0.509 | 0.057 | -0.0026 | -0.0012 | -0.0001 | -0.0012 | - |
| Hip SampEn | 0.115 (0.000) | 0.459 | F(3,57) = 0.98† | 0.359 | 0.509 | 0.049 | 0.0168 | 0.0200 | 0.0191 | 0.0209 | - |
| Knee SampEn | 0.550 (0.075) | - | F(3,54) = 4.21 | 0.009 | **0.030*** | 0.190 | -0.0228 | -0.0179 | -0.0120 | -0.0182 | - |
| Ankle SampEn | 0.554 (0.079) | - | F(3,54) = 2.55 | 0.065 | 0.127 | 0.124 | 0.1032 | 0.0889 | 0.0798 | 0.0837 | - |
| **Joint angles - Pose2Sim method** | | | | | | | | | | | |
| Hip λshort | 0.956 (0.977) | - | F(3,57) = 4.12 | 0.010 | **0.031*** | 0.178 | -0.0221 | -0.0225 | -0.0314 | -0.0242 | 100% - 140% |
| Knee λshort | 0.372 (0.005) | 0.615 | F(3,54) = 0.50† | 0.594 | 0.685 | 0.027 | 0.0096 | 0.0034 | 0.0070 | 0.0150 | - |
| Ankle λshort | 0.817 (0.642) | - | F(3,54) = 5.95 | 0.001 | **0.008*** | 0.248 | -0.2071 | -0.1955 | -0.1560 | -0.1560 | 100% - 140%  120% - 140% |
| Hip ACI | 0.771 (0.467) | - | F(3,57) = 4.58 | 0.006 | **0.022*** | 0.194 | -0.0052 | -0.0074 | -0.0088 | -0.0053 | 100% - 140% |
| Knee ACI | 0.716 (0.348) | - | F(3,54) = 0.59 | 0.625 | 0.703 | 0.032 | 0.0023 | 0.0017 | 0.0006 | 0.0024 | - |
| Ankle ACI | 0.563 (0.087) | - | F(3,54) = 0.18 | 0.906 | 0.927 | 0.010 | -0.0027 | -0.0023 | -0.0023 | -0.0031 | - |
| Hip SampEn | 0.275 (0.000) | 0.564 | F(3,57) = 8.88† | 0.001 | **0.008*** | 0.319 | 0.0046 | 0.0074 | 0.0136 | 0.0061 | 100% - 140%  120% - 140%  140% - SP |
| Knee SampEn | 0.431 (0.015) | 0.713 | F(3,54) = 5.86† | 0.005 | **0.022*** | 0.245 | -0.0315 | -0.0224 | -0.0200 | -0.0269 | 100% - 120%  100% - 140% |
| Ankle SampEn | 0.464 (0.025) | 0.737 | F(3,54) = 3.71† | 0.030 | 0.070 | 0.171 | 0.0826 | 0.0546 | 0.0506 | 0.0673 | - |
| **Trunk accelerations** | | | | | | | | | | | |
| AP trunk acc. λshort | 0.710 (0.267) | - | F(3,60) = 11.50 | < 0.001 | **< 0.001*** | 0.365 | 0.0107 | 0.0268 | 0.0839 | 0.0105 | 100% - 140%  120% - 140%  140% - SP |
| ML trunk acc. λshort | 0.441 (0.009) | 0.667 | F(3,60) = 29.81† | < 0.001 | **< 0.001*** | 0.599 | 0.3452 | 0.3967 | 0.4673 | 0.3343 | 100% - 120%  100% - 140%  120% - 140%  120% - SP  140% - SP |
| VT trunk acc. λshort | 0.459 (0.012) | 0.705 | F(3,60) = 0.71† | 0.505 | 0.651 | 0.034 | 0.1872 | 0.1665 | 0.1813 | 0.1888 | - |
| AP trunk acc. ACI | 0.822 (0.597) | - | F(3,60) = 1.54 | 0.213 | 0.340 | 0.071 | -0.0065 | -0.0050 | -0.0040 | -0.0033 | - |
| ML trunk acc. ACI | 0.574 (0.065) | - | F(3,60) = 4.67 | 0.005 | **0.022*** | 0.189 | -0.0024 | -0.0018 | -0.0002 | 0.0045 | 100% - SP |
| VT trunk acc. ACI | 0.232 (0.000) | 0.539 | F(3,60) = 9.16† | 0.001 | **0.008*** | 0.314 | -0.0063 | -0.0038 | -0.0034 | -0.0120 | 120% - SP  140% - SP |
| AP trunk acc. SampEn | 0.401 (0.004) | 0.668 | F(3,60) = 2.49† | 0.096 | 0.172 | 0.111 | 0.1051 | 0.1155 | 0.1178 | 0.1034 | - |
| ML trunk acc. SampEn | 0.313 (0.001) | 0.565 | F(3,60) = 5.16† | 0.015 | **0.041*** | 0.205 | -0.0989 | -0.0750 | -0.0662 | -0.0894 | - |
| VT trunk acc. SampEn | 0.210 (0.000) | 0.576 | F(3,60) = 4.99† | 0.016 | **0.042*** | 0.200 | -0.0109 | -0.0077 | 0.0248 | -0.0164 | 120%-140% |

**Notes:** †: Greenhouse-Geisser corrected (sphericity violated, Mauchly p < 0.05). εGG: Greenhouse-Geisser epsilon (reported only when correction was applied). p: p-value used for inference (Greenhouse-Geisser corrected when sphericity was violated, uncorrected otherwise). pFDR: p-value after Benjamini-Hochberg false discovery rate correction across all 45 tests; * pFDR < 0.05. η²p: partial eta squared. Bias: mean inter-system difference (markerless minus Vicon) per speed condition; 100%, 120%, 140%: percentage of preferred walking speed. SP: self-paced. Post-hoc pairs: Bonferroni-adjusted pairwise comparisons reaching significance (p < 0.05), reported only for measures with pFDR < 0.05. GC: generalized-coordinate method. P2S: Pose2Sim. λshort: maximum Lyapunov exponent. ACI: attractor complexity index. SampEn: sample entropy. AP: anteroposterior. ML: mediolateral. VT: vertical.

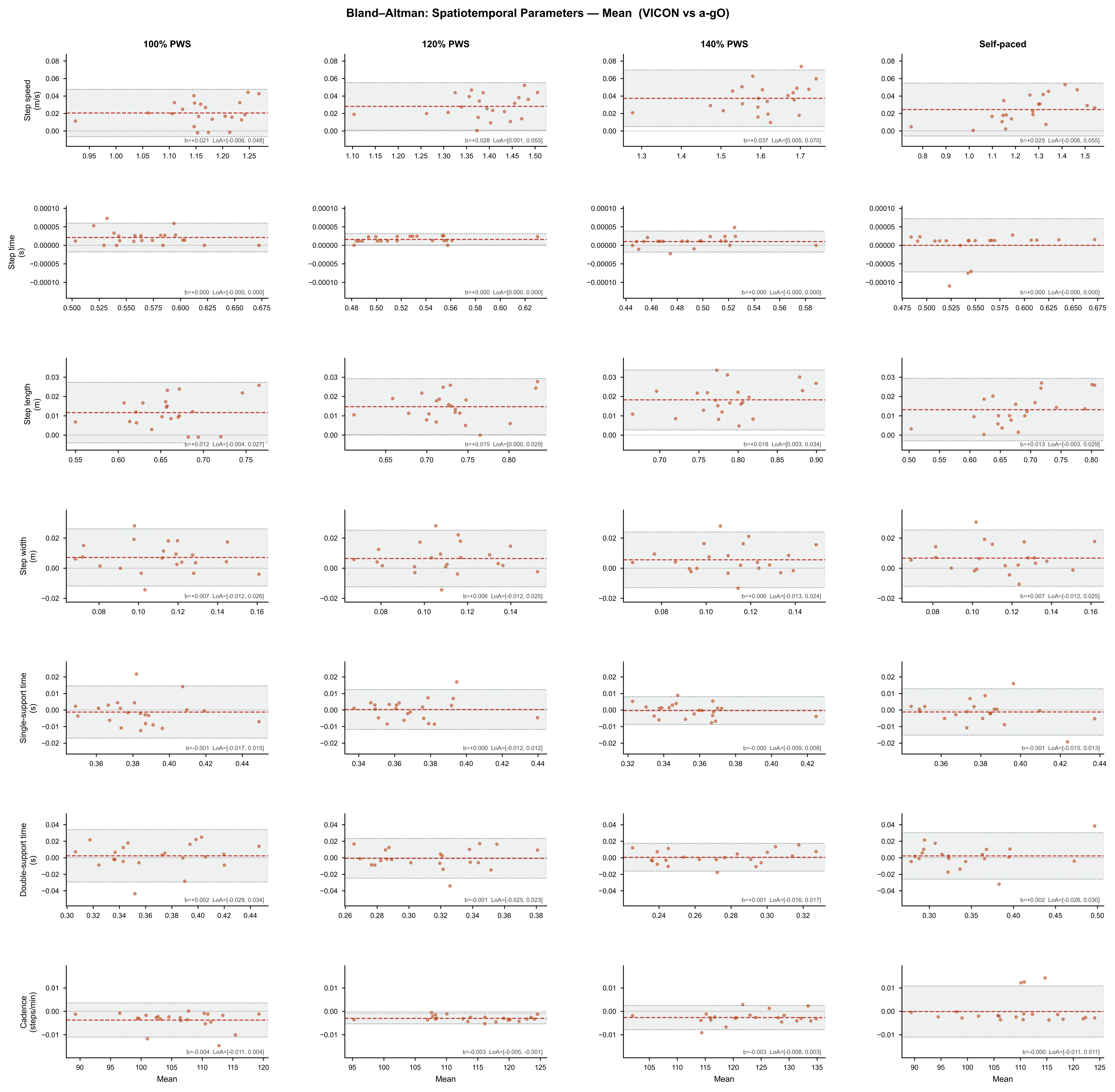
**Supplementary Figure S1**. Bland-Altman for the mean of spatiotemporal parameters across walking conditions.

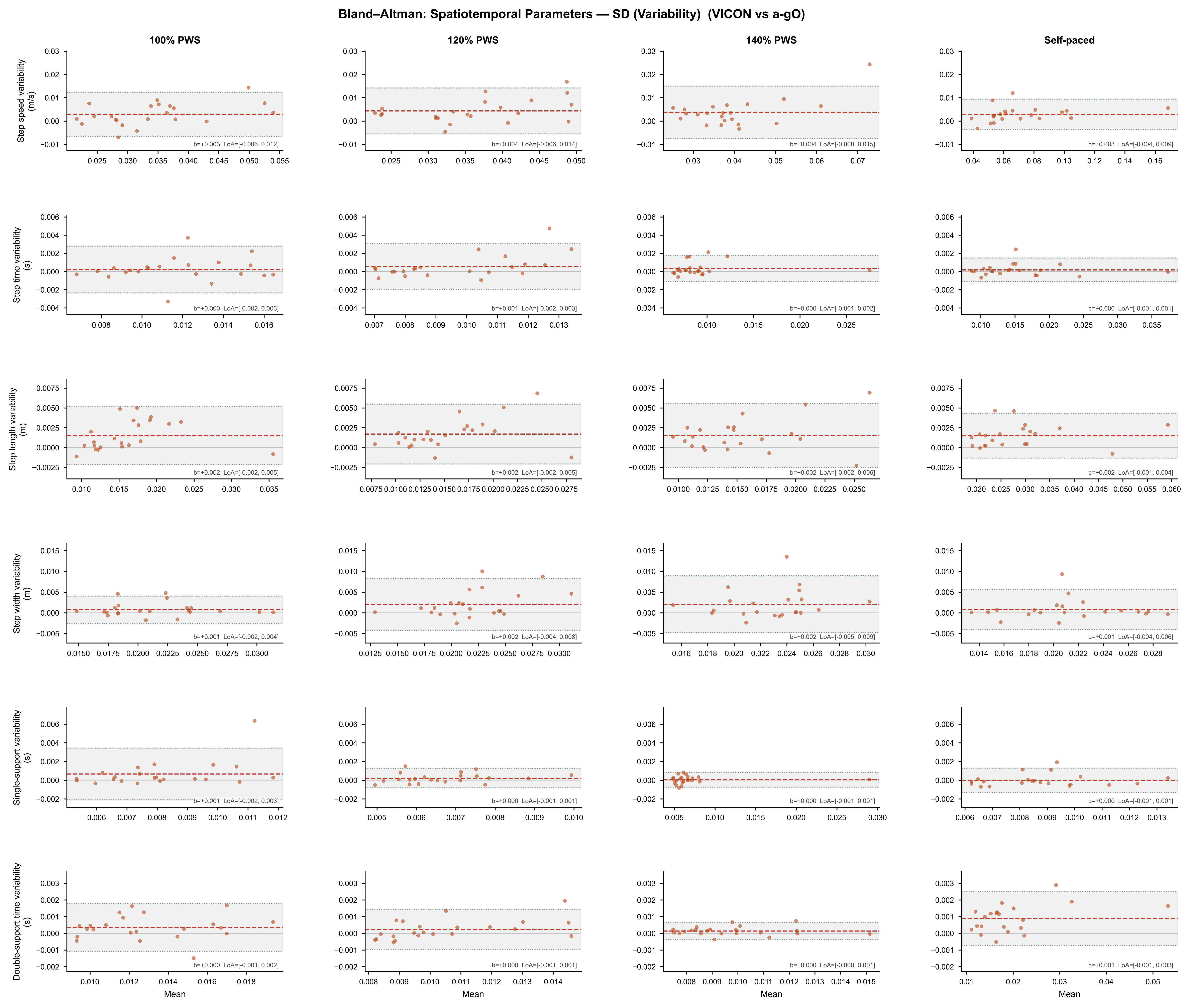
**Supplementary Figure S2**. Bland-Altman for the standard deviation of spatiotemporal parameters across walking conditions.

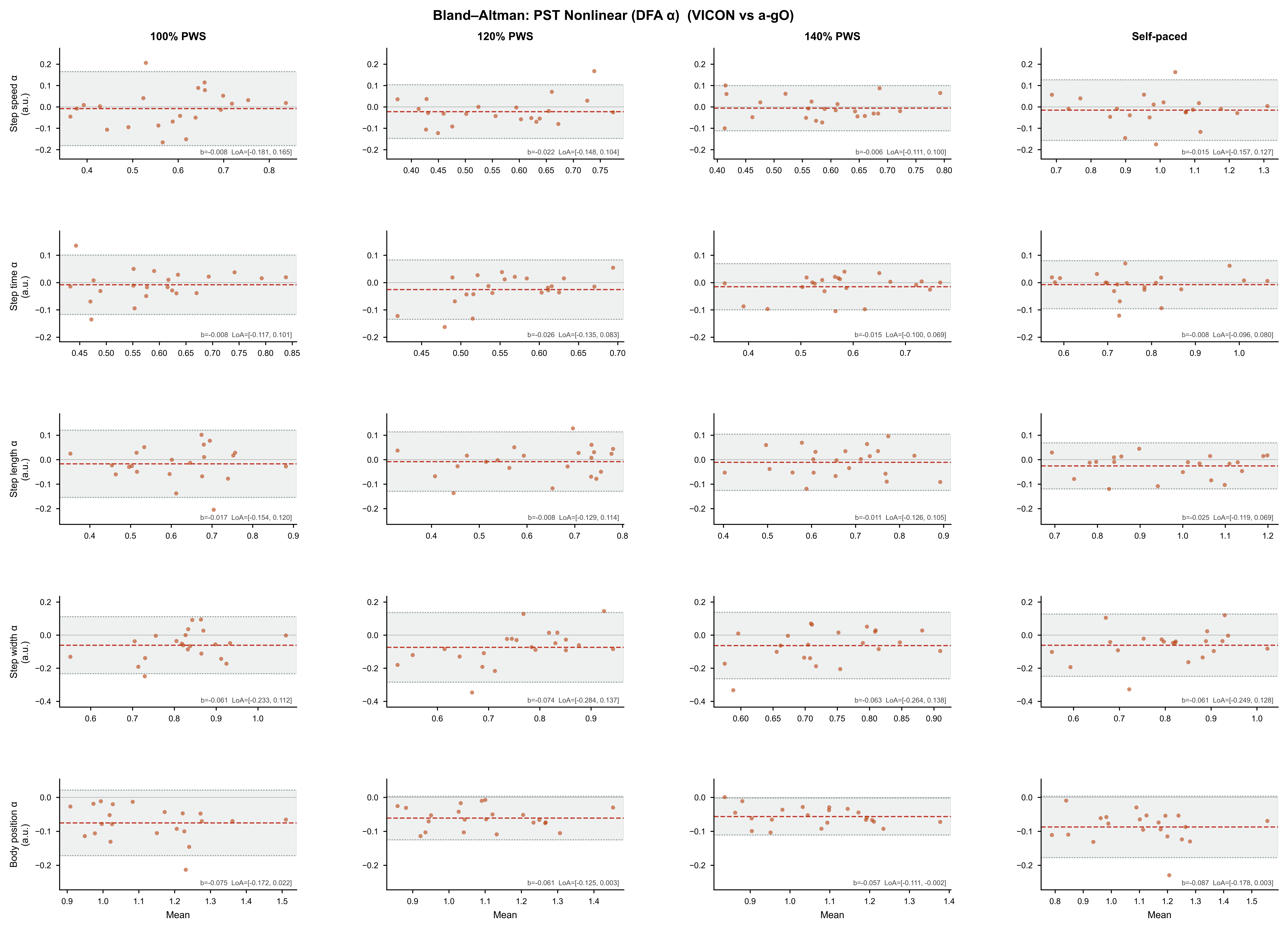
**Supplementary Figure S3**. Bland-Altman for the temporal structure of spatiotemporal parameters across walking conditions.

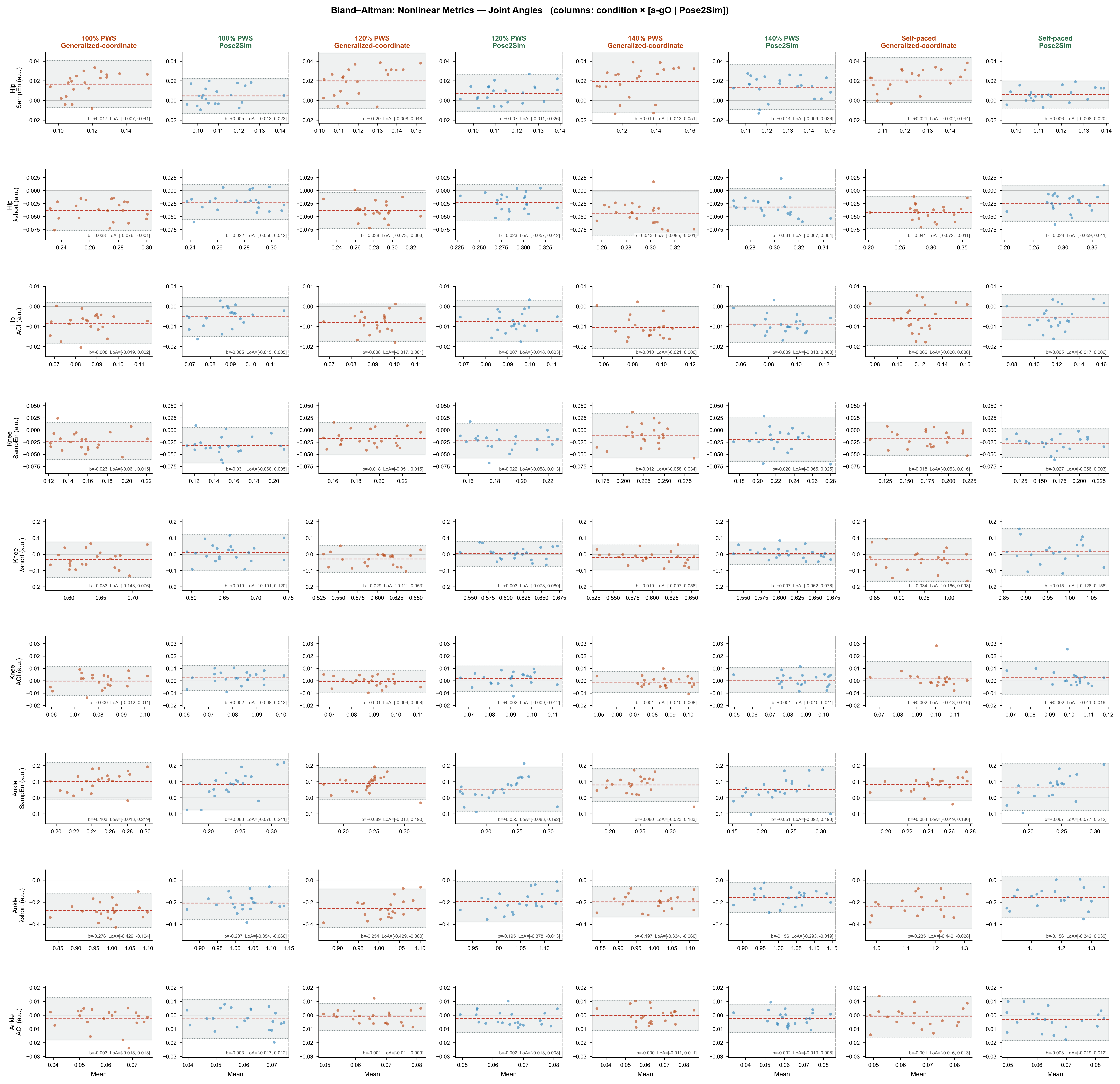
**Supplementary Figure S4**. Bland-Altman for the nonlinear analyses applied on gait joint angles across walking conditions, separately for the generalized-coordinate and Pose2Sim methods.

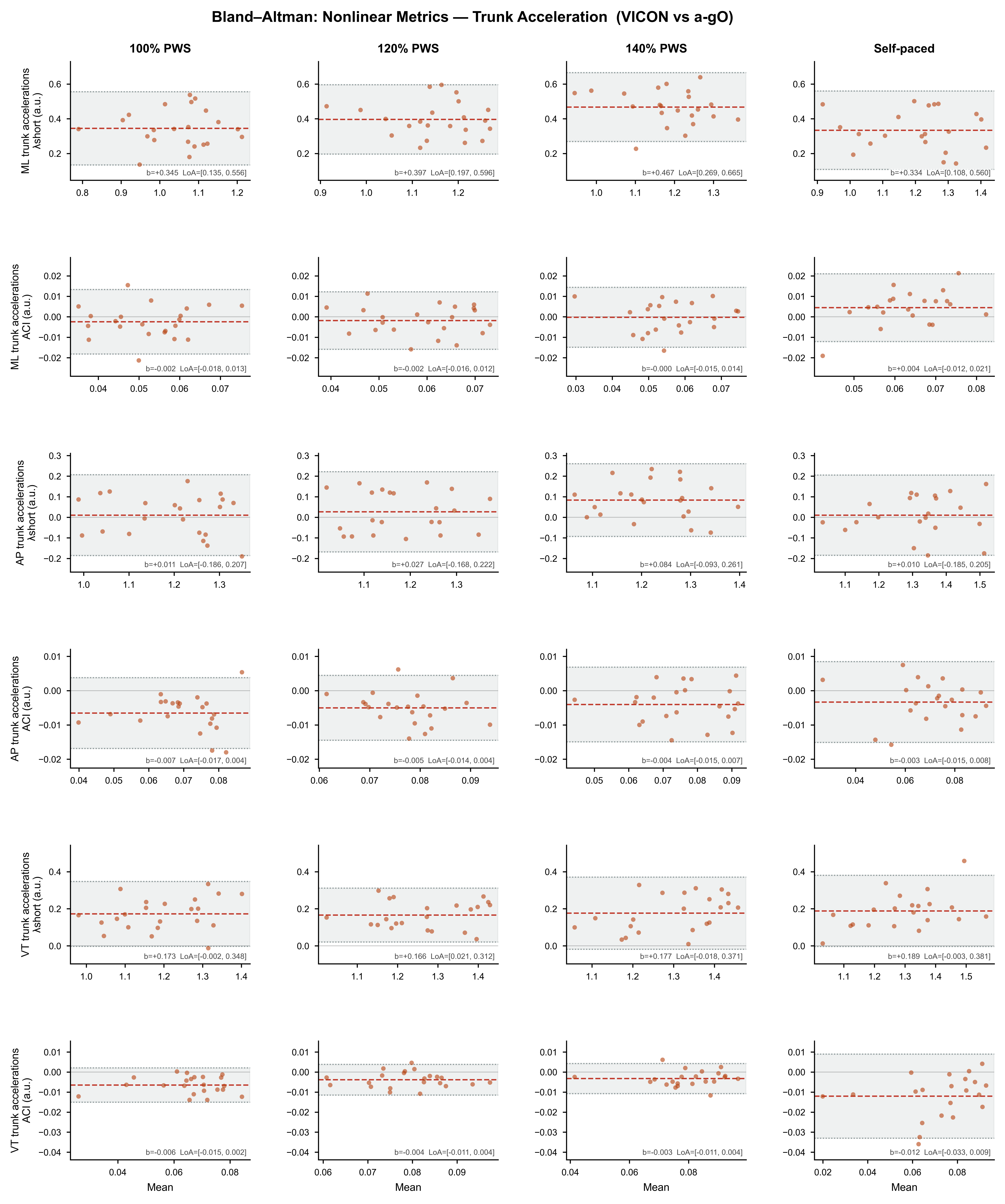
**Supplementary Figure S5**. Bland-Altman for the nonlinear analyses of trunk accelerations across walking conditions.
